# An Enhanced Conditional Variational Autoencoder-Based Normative Model for Neuroimaging Analysis

**DOI:** 10.1101/2025.01.05.631276

**Authors:** Mai P. Ho, Yang Song, Perminder S. Sachdev, Jiyang Jiang, Wei Wen

## Abstract

Normative modelling in neuroimaging provides a powerful framework for quantifying individual deviations from expected brain measures as a function of relevant covariates. While most established approaches have focused on analysing distinct variables in isolation, there is a growing need for deep learning-based methods capable of handling multiple response variables simultaneously. Conditional variational autoencoders (cVAEs) have previously been applied in this context; however, existing inference methods still face challenges in providing reliable probabilistic predictions.

In this study, we introduce an enhanced cVAE-based framework for normative modelling of neuroimaging data. Our key contribution is the development of a novel inference method that integrates latent space sampling and bootstrapping. This method directly leverages the generative nature of cVAEs to estimate the conditional distributions of brain measures. We demonstrate the effectiveness of this approach using white matter hyperintensity (WMH) volumes as a test case for developing, testing, and validating the model. Our dataset includes 8,551 normotensive and 18,180 hypertensive participants from the UK Biobank. We evaluated our method against three well-established and popular normative modelling techniques, including Generalised Additive Models for Location, Scale, and Shape (GAMLSS), Multivariate Fractional Polynomial Regression (MFPR), and Hierarchical Bayesian Regression (HBR).

Our results indicate that the proposed cVAE-based framework achieves comparable performance across various metrics, while capturing individual deviations that correlate with the severity of hypertension. The model is particularly well-suited for high-dimensional, non-linear data, making it a robust tool for assessing individual deviations in brain structure. This enhanced normative modelling architecture paves the way for more nuanced and reliable assessments of brain health, with potential implications for personalised medicine and early detection of neurological disorders.

## 1. Introduction

Normative modelling has recently emerged as a powerful approach in neuroimaging analysis for understanding individual variations in brain structure and function. Unlike traditional case-control studies that focus on group-level differences, normative modelling aims to create a statistical norm of brain measures across populations, against which individual subjects can be compared (Marquand, Rezek, et al., 2016). This approach allows for the quantification of individual deviations from expected brain measures, potentially revealing subtle morphological differences that might go undetected using conventional techniques (Fraza et al., 2021). As a result, normative modelling holds great potential for early detection of neurological disorders and assessment of disease progression.

Several statistical approaches have been employed in normative modelling. Ge et al. (2024) provided a comprehensive comparison of these methods, highlighting widely-used algorithms such as Generalised Additive Models for Location, Scale, and Shape (GAMLSS) (Rigby & Stasinopoulos, 2005), Multivariate Fractional Polynomial Regression (MFPR) (Royston & Altman, 1994) and Hierarchical Bayesian Regression (HBR) (Lindley & Smith, 1972). Each of these methods offers distinct strengths and limitations in the context of neuroimaging analysis. For instance, GAMLSS has proven effective for modelling non-linearity through smoothing functions, as demonstrated in lifespan brain development studies (Bethlehem et al., 2022). However, it is not inherently designed to analyse multiple response variables simultaneously, which is crucial for capturing the complex spatial correlations often present in neuroimaging data (Habeck, 2010). Similarly, while HBR is capable of capturing non-linear relationships using basis functions (de Boer et al., 2024), its computational demands and the approach of analysing data points in isolation present challenges. MFPR, known for its superior accuracy and generalisability in the CentileBrain study (Ge et al., 2024), offers flexibility in modelling non-linear relationships using fractional polynomial terms for multivariate data. Despite this, MFPR may face scalability limitations in high-dimensional datasets, as the model complexity grows rapidly with the number of variables (Royston & Altman, 1994).

The limitations of traditional statistical approaches motivated the exploration of deep learning methods for normative modelling. Deep learning has shown great promise in handling complex, non-linear relationships and interactions between variables simultaneously (LeCun et al., 2015), making it especially useful for analysing the rich, high-dimensional information contained in brain scans. In 2019, Pinaya and colleagues introduced a conditional autoencoder (cAE)-based normative model, designed to learn the underlying neuroimaging data structure of healthy individuals while incorporating age and sex as covariates. Briefly, the cAE architecture consists of an encoder that compresses the input data into a latent representation, and a decoder that reconstructs the original input from this latent representation, both conditioned on the given covariates. Building on this work, Lawry Aguila et al. (2022) proposed a conditional variational autoencoder (cVAE)-based approach, which enables the model to quantify uncertainty through the probabilistic representations in the latent space. These AE-based models marked a significant shift towards more complex, non-linear modelling of brain data, leveraging the capability of deep learning to handle multiple variables simultaneously. However, they also come with certain important assumptions. Specifically, both approaches rely on reconstruction errors – the difference between the original input data and model’s reconstruction – to infer deviations from the normative range (Lawry Aguila et al., 2022; Pinaya et al., 2019). This method assumes that larger reconstruction errors correspond to greater deviations from the norm. However, a key limitation lies in how the reconstruction is generated. The model uses both new covariates and the observed brain data to produce the reconstruction, while ideally, predictions should be generated solely from covariates to assess meaningful deviations. This dual input nature of the reconstruction process makes it challenging to determine whether the observed deviations are due to genuine abnormalities in brain structure. Furthermore, these models do not provide probabilistic predictions, which are particularly valuable in normative modelling for understanding individual variations in brain structure and function across a population. As a result, these limitations can affect the interpretability and reliability of conclusions drawn from such models.

In this study, we propose an alternative approach for obtaining predictions from a cVAE-based normative model. Instead of relying on reconstruction errors, we leverage the generative capabilities of cVAEs and employ bootstrapping to directly model the distribution of multiple brain measures conditioned on relevant covariates. To demonstrate our model’s performance on neuroimaging data, white matter hyperintensity (WMH) lesions caused by cerebral small vessel diseases across different brain regions (see Table S1) were used as an example. We modelled WMH volumes by incorporating relevant covariates, including age, sex, total intracranial volume and several vascular risk factors. Age and sex are fundamental factors in brain morphology’s variations, while total intracranial volume accounts for head size. The modifiable vascular risk factors we included – diabetes, hypercholesterolemia, obesity, and smoking – are considered key contributors to cerebrovascular health and are associated with the manifestation of WMH on brain scans. Finally, to demonstrate the clinical usage of the proposed model, we examine the relationship between individual deviations from normative WMH volumes and hypertension levels, which have been shown to correlate positively with white matter burden in the brain (de Leeuw et al., 2002; Du et al., 2024). In addition, we trained normative model using GAMLSS, MFPR and HBR algorithms using the same data for comparison with our cVAE-based approach. Through these analyses, we aim to demonstrate that our proposed cVAE-based normative model can overcome the drawbacks outlined for both traditional statistical and deep learning-based methods while maintaining comparable performance. Our goal is to equip researchers and clinicians with more accurate, interpretable tools for understanding individual brain variations across both healthy and diseased populations.

## 2. Methods and Materials

### 2.1 Participants

The data for this study were obtained from UK Biobank (project #98013). UK Biobank is a large-scale cohort study that includes comprehensive health information from over half a million UK participants (Sudlow et al., 2015). A flowchart detailing the selection of participants for the current study is presented in Figure 1. Initially, 48,458 participants with both T1-weighted (T1w) and T2-weighted-Fluid-Attenuated Inversion Recovery (T2w-FLAIR) scans were considered. Several exclusion criteria were applied as follows: participants with incompatible or unusable T1w/FLAIR scans (n = 3,457), those with intracranial volume z-scores greater than 3 (n = 574), individuals diagnosed with specific neurological disorders outlined in Table S2 (n = 2,772), and those with incomplete data on required variables to determine vascular risk factors, e.g. diabetes diagnosis by doctor (n = 4,295). Additionally, to eliminate potential confounding effects from antihypertensive use on the study’s outcomes, 10,629 participants who were on antihypertensive medication were excluded. After these exclusions, a total of 26,731 participants remained for analysis.

**Figure 1.**
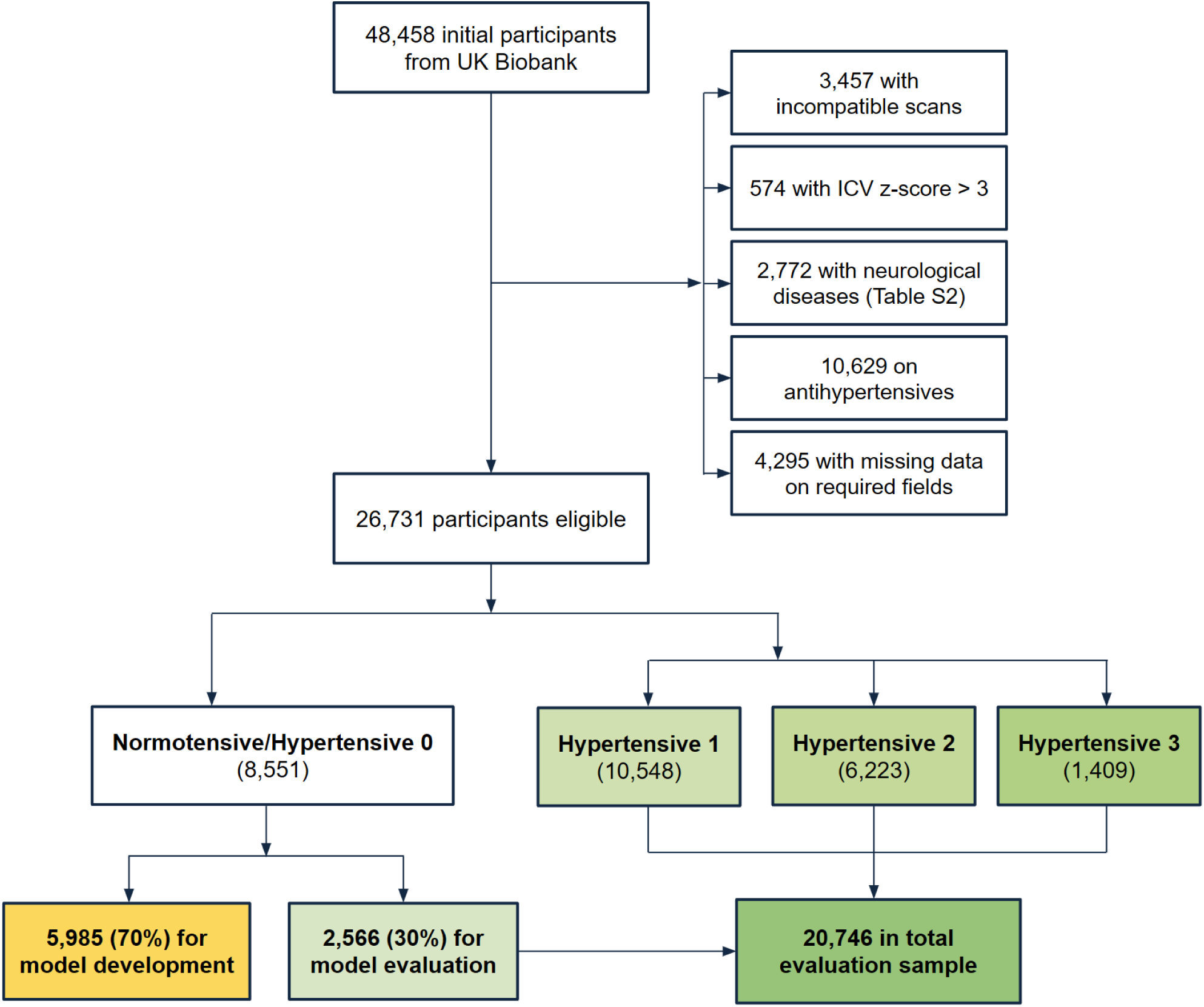
Flowchart of Participant Selection. Abbreviations: T1w = T1-weighted; T2w-FLAIR = T2-weighted-Fluid-Attenuated Inversion Recovery; ICV = Intracranial Volume; SBP = Systolic Blood Pressure; DBP = Diastolic Blood Pressure.

For clinical relevance assessment, the eligible participants were classified into four groups based on blood pressure measurements, following the Global Hypertension Practice guidelines by the International Society of Hypertension (Unger et al., 2020). The normotensive group (Hypertension 0) comprised 8,551 participants with systolic blood pressure (SBP) < 130 mmHg and diastolic blood pressure (DBP) < 85 mmHg. The hypertensive groups were defined as follows: Hypertensive 1 (n = 10,548) with SBP between 130 and 139 mmHg, or DBP between 85 and 89 mmHg; Hypertensive 2 (n = 6,223) with SBP between 140 and 159 mmHg, or DBP between 90 and 99 mmHg; and Hypertensive 3 (n = 1,409) with SBP ≥ 160 mmHg or DBP ≥ 100 mmHg.

For model development, 70% (n = 5,985) of the normotensive group was randomly selected to form the training dataset. The remaining 30% (n = 2,566) of the normotensive group served as hold-out dataset, which was used to assess the model’s performance. This hold-out dataset was then combined with all hypertensive participants to create an evaluation sample of 20,746 participants, used to test the model’s clinical relevance across different levels of hypertension severity.

### 2.2 Vascular Risk Factors

Several vascular risk factors other than hypertension were considered as part of the covariates. These factors, treated as binary covariates, were diabetes, hypercholesterolemia, obesity and smoking. Participants with diabetes were identified based on a doctor’s diagnosis and the use of antidiabetic medication. Hypercholesterolemia was determined through the participant medication records, indicating treatment for elevated cholesterol levels. Obesity was defined as having a body mass index (BMI) of 30 or higher. Smoking status was identified as current or former smoker based on participants’ self-reported smoking history. Controlling for these factors allowed us to better assess the relationship between WMH volumes and hypertension, minimising the influence of other vascular conditions on the results.

### 2.3 MRI Acquisition and Data Preprocessing

T1w and FLAIR scans were acquired from three UK imaging centres (Cheadle Greater Manchester, Newcastle and Reading), all using a 3T Siemens Skyra scanner with a standard Siemens 32-channel head coil and uniform imaging parameters (Miller et al., 2016). Full details of the imaging protocols are available in the online UK Biobank brain imaging documentation (Smith et al., 2024). Whole-brain intracranial volumes, essential for adjusting for head size, were estimated using the UK Biobank’s imaging-processing pipeline (Alfaro-Almagro et al., 2018).

White matter hyperintensity (WMH) volumes were computed in native space using the UBO pipeline (Jiang et al., 2018), with volumes extracted for different regions of interest. All extracted measures underwent quality control to ensure data integrity. The full list of brain regions and their corresponding abbreviations are presented in Table S1. Three parcellation schemes were applied: (1) whole-brain, periventricular, and deep WMH; (2) lobar WMH volumes, with the brain divided into 11 lobar regions; and (3) WMH volumes based on arterial territories, with parcellation according to 16 blood supply areas (Wen & Sachdev, 2004). As the WMH volumes tend to be highly skewed, we applied a log transformation and Z-score standardisation to normalise the data for subsequent analyses. The normalised WMH volumes for each brain region served as inputs for the development and evaluation of all models.

### 2.4 cVAE-based Model Development

#### 2.4.1 Model Architecture

The model employed in this study is a conditional Variational Autoencoder (cVAE), building on the auto-encoding variational Bayes algorithm introduced by Kingma and Welling (2013). The cVAE architecture consists of two main components: an encoder and a decoder. The encoder maps the input data to a lower-dimensional latent space, while the decoder reconstructs the original data from this latent representation conditioned on covariates. The parameters defining the model architecture, including the dimensionality of latent and hidden layers, were explored using the Optuna framework to identify optimal configurations (Akiba et al., 2019). In this study, the cVAE framework is designed to predict WMH volumes across different brain regions while accounting for covariates, including age, sex, total intracranial volume, and vascular risk factors (diabetes, hypercholesterolemia, obesity and smoking).

The comprehensive cVAE framework is illustrated in Figure 2, highlighting both the training and inference phases. During training, the encoder receives two inputs: (1) the WMH volumes (denoted as 𝑌) comprising 30 features detailed in Table S1, and (2) a set of 7 covariates (denoted as 𝑋). The 7 covariates include age, total intracranial volume, and five binary indicators (sex, diabetes, hypercholesterolemia, obesity, and smoking status), which are transformed into a 12-dimensional vector: one continuous value for age, one for total intracranial volume, and five binary values for the other factors. These inputs pass through a 128-unit hidden layer with rectified linear unit (ReLU) activation, allowing the model to capture non-linear relationships between the WMH volumes and covariates.

**Figure 2.**
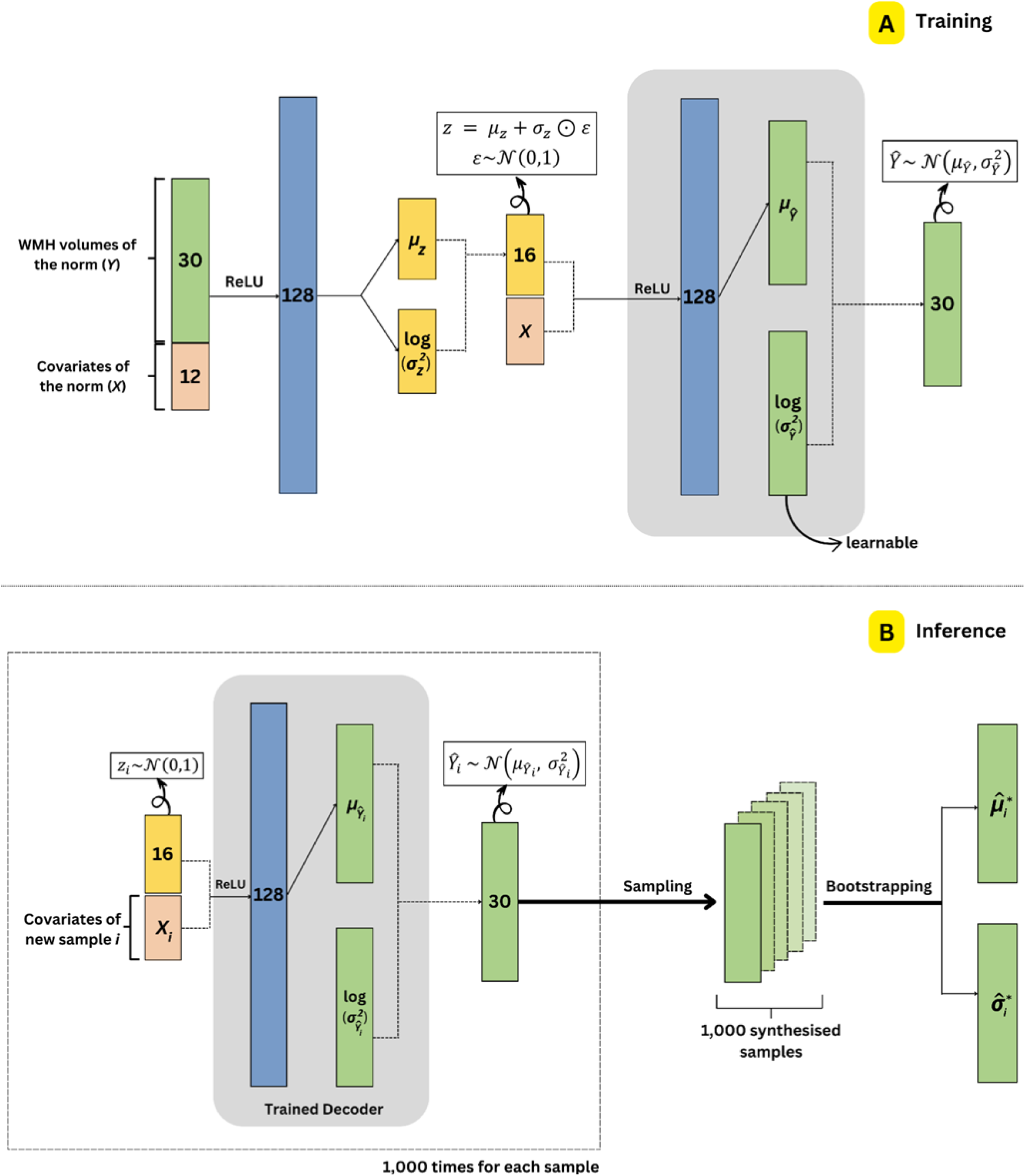
Model Architecture and Inference Framework. This diagram illustrates a conditional Variational Autoencoder (cVAE) architecture for modelling WMH volumes across brain regions. (A) The top panel depicts the model architecture and training phase, showing how 30 WMH features and covariates (transformed into a 12-length vector) are processed through the encoder, latent space and decoder. (B) The bottom panel demonstrates the inference process, where the trained decoder is used to generate new predictions. This process is repeated 1,000 times to synthesise samples, from which summary statistics (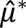 and 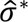) are computed via bootstrapping. Layers represented in the same colour indicate identical dimensions. Abbreviations: WMH = White Matter Hyperintensity; ReLU = Rectified Linear Unit. ⊙ denotes element-wise multiplication.

The encoder outputs two key latent variables: the mean (𝜇_𝑧_) and the log-variance 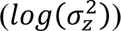, both represented as 16-dimensional vectors. These latent variables define a multivariate Gaussian distribution from which a latent variable 𝑧 is sampled. As proposed by Kingma and Welling (2013), the reparameterization trick is applied to enable backpropagation through stochastic layers. Specifically, z is calculated as:

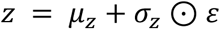

Here, 𝜀∼𝒩(0,1) is a random noise variable sampled from a standard normal distribution, and ⊙ denotes element-wise multiplication. This technique allows the sampling process to be expressed as a deterministic operation, making it possible to compute gradients and update model parameters during training.

Subsequently, the decoder reconstructs the WMH volume data by processing a 28-length input (latent vector 𝑧𝑧 concatenated with covariates) through a 128-unit hidden layer with ReLU activation. The decoder produces two outputs that define a probabilistic distribution over the reconstructed WMH volumes. The first output is the mean vector 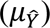, obtained by transforming the 128-dimensional hidden presentation through a linear layer to produce a 30-dimensional vector. The second output is the log-variance 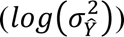, which is implemented as a single global learnable parameter shared across all features. This global variance parameter is updated during training to optimise reconstruction quality. Together, these components define the final output, which is a multivariate normal distribution 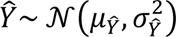 From this distribution, multiple predictions of WMH volumes can be sampled. The rationale behind outputting a distribution rather than a deterministic value is twofold. First, brain structure measurements like WMH volumes vary significantly between individuals (Habes et al., 2016). Modelling the output as a distribution thus better captures this inherent biological variability, which a single point estimate might overlook. Second, a distributional output provides a measure of the model’s confidence in its predictions. This is crucial in clinical applications, where understanding the reliability of a prediction can inform clinical decision-making.

#### 2.4.2 Training Phase

The cVAE is trained by minimising a loss function that combines two terms: the Kullback-Leibler (KL) divergence (Kullback & Leibler, 1951) and the reconstruction loss. The reconstruction loss measures how well the model can reconstruct the original WMH volumes from the latent space, while the KL divergence serves as a regularization term, encouraging the latent space to remain close to a predefined prior distribution (𝒩(0,1)). The overall loss function (𝓛𝓛) of the cVAE is defined as:

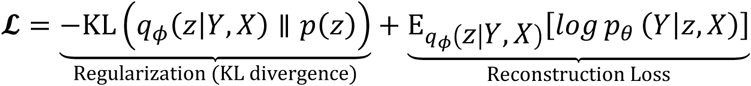

In this equation, the first term is the KL divergence, which measures the difference between the learned posterior distribution 𝑞_𝜙_(𝑧 | 𝑌, 𝑋), and the Gaussian prior distribution 𝑝(𝑧). The second term represents the expected log-likelihood of generating the input 𝑌 given the latent representation 𝑧𝑧 and covariates 𝑋. The encoder and decoder parameters are denoted as 𝜙 and 𝜃, respectively.

The model is trained using the Adam optimiser, with hyperparameter tuning performed using the Optuna framework (Akiba et al., 2019). To prevent overfitting, early stopping is implemented with a patience of 20 epochs.

#### 2.4.3 Inference Phase

The inference phase of this framework leverages the cVAE’s generative capabilities to produce robust probabilistic predictions for each sample in the evaluation dataset. Specifically, for each set of input covariates 𝑋_𝑖_ for participant 𝑖, a 16-dimensional latent vector 𝑧_𝑖_ is sampled from the standard normal distribution 𝒩(0,1), which is the predefined prior distribution for the latent space. This sampled vector is combined with the input covariates 𝑋_𝑖_to form a concatenated input, which is then passed through the trained cVAE decoder to output a predicted distribution 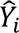 for the WMH volumes. The predicted distribution is parameterised as 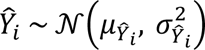, where 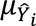 is the mean and 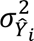 is the variance derived from the log-variance output by the decoder. From this predicted distribution, a single sample is drawn to represent one possible reconstruction for the WMH volumes given the covariates 𝑋_𝑖_. To adequately capture the inherent variability in the data, this process is repeated 1,000 times for each set of covariates. This results in 1,000 synthesised samples per input, providing a comprehensive range of plausible reconstructions of the WMH volumes.

To further enhance the robustness of these predictions, a bootstrapping method is applied to the synthesised samples. Specifically, 1,000 bootstrap iterations are conducted, where each iteration involves randomly sampling with replacement from the set of synthesised outcomes for each sample in the evaluation dataset. For each bootstrap sample, summary statistics – such as the mean and standard deviation of the predicted WMH volumes – are calculated. The averages of these summary statistics across all bootstrap iterations are then used to produce the final probabilistic predictions (e.g., aggregated mean 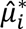, and standard deviation 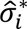) for each participant 𝑖.

This inference framework accounts for both the model’s inherent variability and the uncertainty associated with finite sampling. Such an approach is particularly valuable in clinical applications, where understanding the range of possible outcomes and their associated uncertainties is crucial for informed decision-making and interpretation of results.

### 2.5 Comparable Models

We compared the cVAE with three well-established models that are commonly used for normative modelling: Generalised Additive Models for Location, Scale, and Shape (GAMLSS), Multivariate Fractional Polynomial Regression (MFPR), and Hierarchical Bayesian Regression (HBR).

For the GAMLSS model, spline functions were incorporated to capture the non-linear relationships between predictors and outcomes. In the case of MFPR, non-linearity was addressed by systematically selecting the most appropriate fractional polynomial functions to model the predictor-outcome relationship. The HBR model, implemented using the PCNtoolkit package (Kia et al., 2020), handles non-linearity through its hierarchical structure, which incorporates non-linear basis functions. All comparable models were trained using 5-fold cross-validation to ensure robust performance estimation and mitigate overfitting.

### 2.6 Deviation Score Computation

To quantify the deviation of observed values from model predictions, a standardised deviation score was computed. Specifically, the deviation score 𝑧_𝑖*j*_ for participant 𝑖 and feature 𝑗 is defined as:

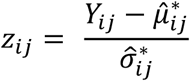

where 𝑌_𝑖_*_j_* is the observed value; 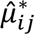 denotes the predicted mean; and 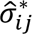 represents the predicted standard deviation for participant 𝑖 and feature 𝑗.

### 2.7 Model Performance Metrics

Despite the normalisation of input WMH volumes during data preprocessing, both the input and synthesised data exhibited non-Gaussian characteristics. As a result, we selected a set of performance metrics that are robust to non-Gaussian distributions to compare the performance of the cVAE against other models. These metrics include (1) *Median Absolute Error*, (2) *Quantile Loss at the 90^th^ Percentile*, (3) *Spearman’s Correlation Coefficient*, and (4) *Explained Variance*.

*Median Absolute Error* was used to measure the typical prediction error across all participants by calculating the median of the absolute differences between observed and predicted mean values. Unlike Mean Absolute Error, it is less influenced by outliers, making it more suitable for non-Gaussian data. *Quantile Loss at the 90th Percentile* assesses the model’s accuracy in predicting extreme values by focusing on the upper end (90^th^ percentile) of the error distribution. This highlights the model’s ability to capture the upper tail, essential for identifying extreme cases. *Spearman’s Correlation Coefficient* was employed to assess the monotonic relationship between observed and predicted values without assuming linearity. Meanwhile, *Explained Variance*, a common metric in normative modelling studies (Ge et al., 2024; Rutherford et al., 2022), indicates the proportion of variability in observed values that each model can capture. The mathematical formulation and interpretation of each metric are provided in Table 1.

**Table 1.**
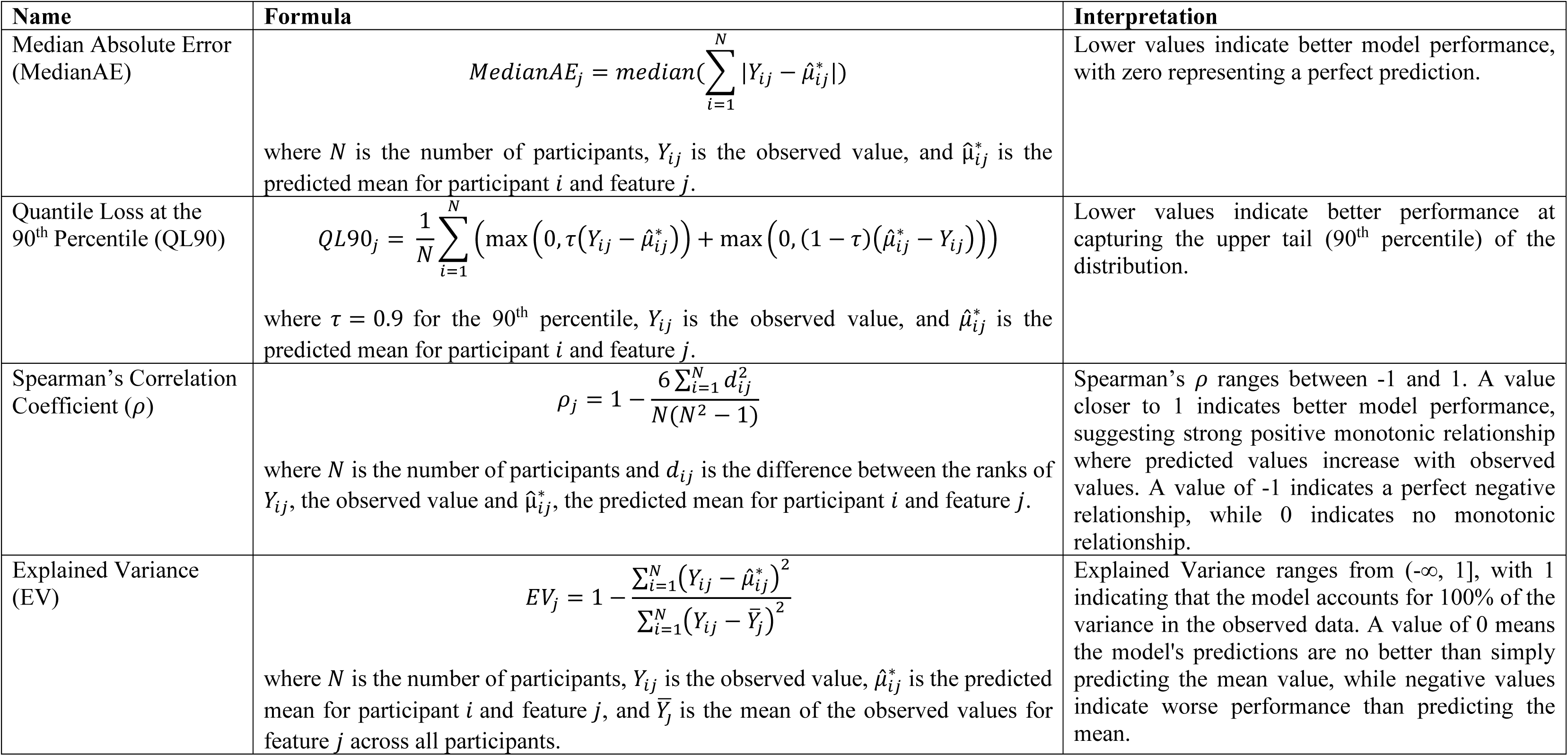
Performance metrics.

To assess whether the performance differences between the cVAE model and each comparison model (GAMLSS, MFPR, and HBR) were statistically significant, median-based permutation tests were conducted. This non-parametric approach evaluates if observed differences in performance metrics could have occurred by chance. Briefly, for each feature, the observed median difference in performance (e.g., Median Absolute Error) between the cVAE model and a given comparison model was first calculated. Next, a null distribution was constructed by pooling all feature-wise scores from both models and randomly partitioning them into two groups of their original sizes. The median difference between the two reshuffled groups was then computed, and this procedure was repeated 10,000 times to form a distribution of median differences expected under the null hypothesis of no true difference between models. The empirical p-value was calculated as the proportion of shuffled differences with an absolute value greater than or equal to that of the observed difference. This process was repeated independently for each model comparison, ensuring a robust, distribution-free assessment of statistical significance across non-Gaussian data.

### 2.8 Deviation Analysis

#### 2.8.1 Deviation Score Analysis on Hold-Out Dataset

On the hold-out dataset, z-scores were computed to assess how individual data points deviate from the normative model predictions. First, we examined the relationship between age and z-scores to evaluate the model’s ability to generate predictions that were independent of covariates, particularly age. This analysis was essential in ensuring that the deviation scores reflected clinically relevant abnormalities, rather than being confounded by age or other covariates.

Next, we analysed the percentage of samples with z-scores greater than 2.58, corresponding to a 99% confidence interval. Extreme deviations from the normative range have been shown to correlate with clinical symptoms in individual patients (Fraza et al., 2021; Marquand, Wolfers, et al., 2016). Given that the hold-out dataset shares similar characteristics with the training data, lower rates of extreme deviations are expected and indicate that the model has accurately captured the typical range of variations seen in the training set. Conversely, a higher percentage of extreme deviations could suggest overfitting, potentially resulting in the misclassification of normal variations as abnormalities.

#### 2.8.2 Clinical Application

In the entire evaluation dataset, we explored the relationship between WMH volume z-scores and different hypertension levels (graded from 0 to 3) using the Spearman’s correlation test. Additionally, the percentage of extreme deviations (z-score > 2.58) was calculated and compared across the different hypertension levels to understand how hypertension influences WMH abnormalities.

## 3. Results

### 3.1 Sample Characteristics and WMH Volume Distribution

Sample characteristics, including demographics and vascular risk factors of train and evaluation data, are summarised in Table 2. The training sample consisted of 5,985 normotensive individuals (referred to as hypertension level 0 in Table 2), with a mean age of 60.84 ± 7.20 years, of whom 30.8% were male. The evaluation sample included 20,746 participants, further subdivided into 2,566 normotensive (mean age 60.90 ± 7.31 years, 30.8% male) and 18,180 hypertensive individuals (mean age 64.69 ± 7.49 years, 48.6% male). Hypertension levels were classified into three groups (1-3), with 58.0% of hypertensive individuals falling into level 1, 34.2% into level 2, and 7.8% into level 3. Hypertensive participants exhibited higher rates of vascular risk factors, including diabetes, hypercholesterolemia, obesity, and smoking, compared to normotensive participants.

**Table 2.**
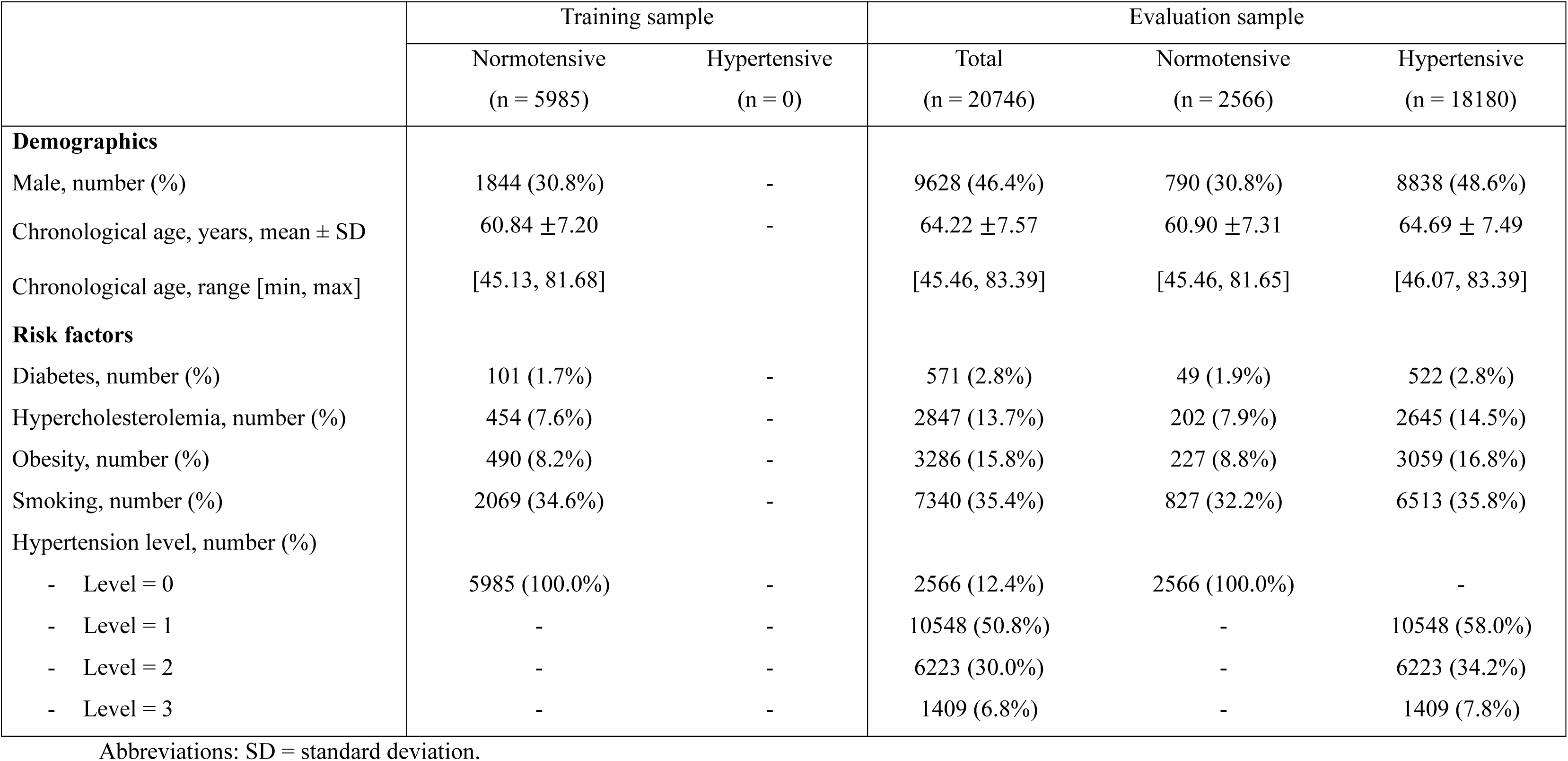
Characteristics of samples (n = 26,731).

|  | Training sample |  | Evaluation sample |  |  |
| --- | --- | --- | --- | --- | --- |
|  | Normotensive<br>(n = 5985) | Hypertensive<br>(n = 0) | Total<br>(n = 20746) | Normotensive<br>(n = 2566) | Hypertensive<br>(n = 18180) |
| **Demographics** |  |  |  |  |  |
| Male, number (%) | 1844 (30.8%) | - | 9628 (46.4%) | 790 (30.8%) | 8838 (48.6%) |
| Chronological age, years, mean ± SD | 60.84 ± 7.20 | - | 64.22 ± 7.57 | 60.90 ± 7.31 | 64.69 ± 7.49 |
| Chronological age, range [min, max] | [45.13, 81.68] |  | [45.46, 83.39] | [45.46, 81.65] | [46.07, 83.39] |
| **Risk factors** |  |  |  |  |  |
| Diabetes, number (%) | 101 (1.7%) | - | 571 (2.8%) | 49 (1.9%) | 522 (2.8%) |
| Hypercholesterolemia, number (%) | 454 (7.6%) | - | 2847 (13.7%) | 202 (7.9%) | 2645 (14.5%) |
| Obesity, number (%) | 490 (8.2%) | - | 3286 (15.8%) | 227 (8.8%) | 3059 (16.8%) |
| Smoking, number (%) | 2069 (34.6%) | - | 7340 (35.4%) | 827 (32.2%) | 6513 (35.8%) |
| Hypertension level, number (%) |  |  |  |  |  |
| - Level = 0 | 5985 (100.0%) | - | 2566 (12.4%) | 2566 (100.0%) | - |
| - Level = 1 | - | - | 10548 (50.8%) | - | 10548 (58.0%) |
| - Level = 2 | - | - | 6223 (30.0%) | - | 6223 (34.2%) |
| - Level = 3 | - | - | 1409 (6.8%) | - | 1409 (7.8%) |

Abbreviations: SD = standard deviation.

Table S3 provides descriptive statistics of WMH volumes (in mm³) across various brain regions in both the training and evaluation samples. In the evaluation dataset, WMH volumes were consistently higher across brain regions in hypertensive individuals than in normotensive participants. For example, whole-brain WMH volumes increased from 1761.5 ± 3241.66 mm³ in normotensive participants to 2433.40 ± 3315.97 mm³ in those with Hypertensive Level 1, eventually reaching 3933.86 ± 5122.58 mm³ in Hypertensive Level 3. However, certain regions, such as the cerebellum and posterior artery callosal (PAC) areas, consistently displayed lower WMH volumes. Notably, the differences between normotensive and hypertensive groups in these regions were relatively small. For instance, the right cerebellum WMH volume was 0.33 ± 3.35 mm³ in normotensive participants compared to 0.35 ± 3.42 mm³ in those with Hypertensive Level 3.

Figure S1 displays the distribution of whole-brain WMH volumes before and after concurrent log transformation and z-score normalisation.

### 3.2 Correlations between Covariates and WMH Volumes

The Spearman and point-biserial correlations between covariates and region-specific WMH volumes in the training dataset are illustrated in Figure S2 and detailed in Table S4. As shown in Figure S2, age showed significant positive correlations with WMH volumes across multiple brain regions in the training dataset. According to Table S4, the strongest correlations with age were observed in general measures, including whole-brain (r = 0.398, p < 0.001), periventricular (r = 0.374, p < 0.001), and deep white matter (r = 0.355, p < 0.001) regions. In contrast, region such as cerebellum (left: r = −0.033, p = 0.012; right: r = −0.020, p = 0.120) and posterior artery thalamic and midbrain perforators (PATMP) (left: r = −0.017, p = 0.191; right: r = 0.038, p = 0.003) exhibited weak or non-significant correlations with age. Other covariates, including sex and vascular risk factors, demonstrated low to moderate correlations with WMH volumes in all brain regions.

### 3.3 Model Performance Metrics Comparison

Table 3 presents the results of the median-based permutation tests, comparing the performance of cVAE against GAMLSS, MFPR, and HBR models using the following metrics: Median Absolute Error, Quantile Loss at the 90^th^ percentile, Spearman’s Correlation Coefficient between observed and predicted values, and Explained Variance. Detailed performance metric values are provided in Table S5.

**Table 3.**
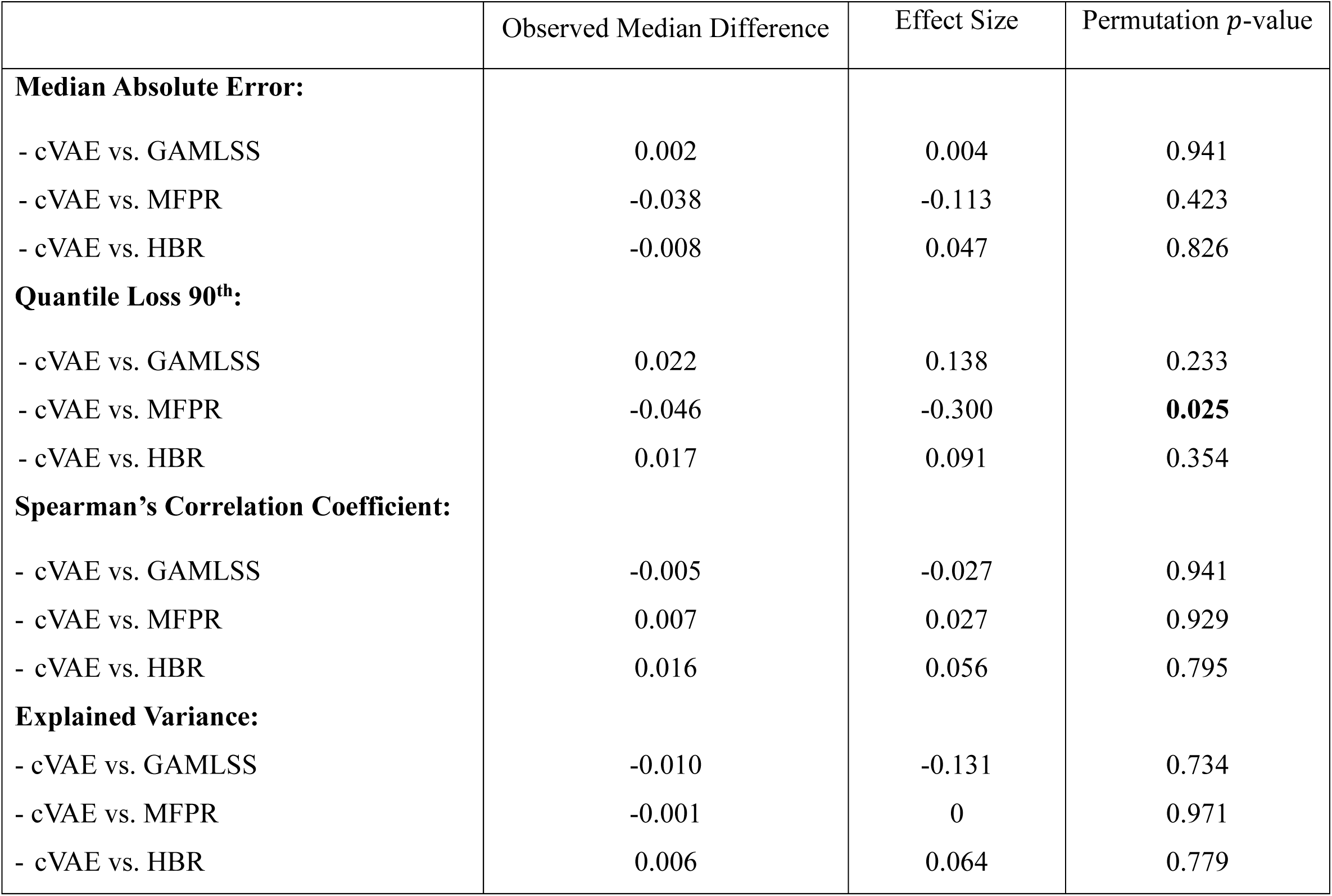
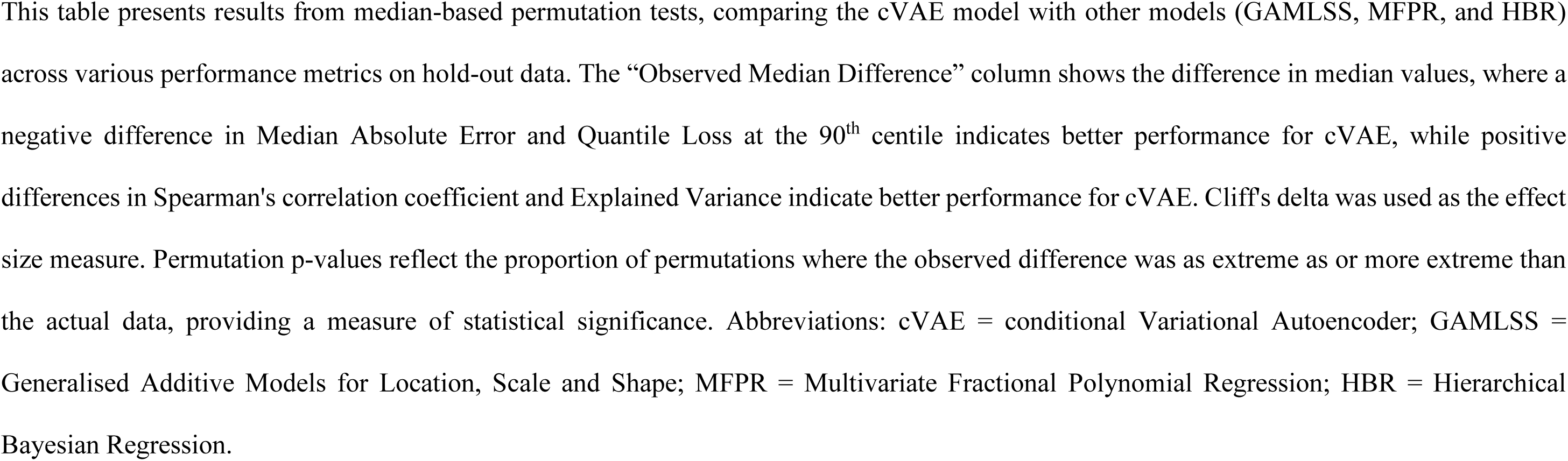
Median-based permutation test results for comparing model performance across metrics on hold-out dataset.

|  | Observed Median Difference | Effect Size | Permutation *p*-value |
| --- | --- | --- | --- |
| **Median Absolute Error:** |  |  |  |
| - cVAE vs. GAMLSS | 0.002 | 0.004 | 0.941 |
| - cVAE vs. MFPR | -0.038 | -0.113 | 0.423 |
| - cVAE vs. HBR | -0.008 | 0.047 | 0.826 |
| **Quantile Loss 90<sup>th</sup>:** |  |  |  |
| - cVAE vs. GAMLSS | 0.022 | 0.138 | 0.233 |
| - cVAE vs. MFPR | -0.046 | -0.300 | **0.025** |
| - cVAE vs. HBR | 0.017 | 0.091 | 0.354 |
| **Spearman's Correlation Coefficient:** |  |  |  |
| - cVAE vs. GAMLSS | -0.005 | -0.027 | 0.941 |
| - cVAE vs. MFPR | 0.007 | 0.027 | 0.929 |
| - cVAE vs. HBR | 0.016 | 0.056 | 0.795 |
| **Explained Variance:** |  |  |  |
| - cVAE vs. GAMLSS | -0.010 | -0.131 | 0.734 |
| - cVAE vs. MFPR | -0.001 | 0 | 0.971 |
| - cVAE vs. HBR | 0.006 | 0.064 | 0.779 |

This table presents results from median-based permutation tests, comparing the cVAE model with other models (GAMLSS, MFPR, and HBR) across various performance metrics on hold-out data. The “Observed Median Difference” column shows the difference in median values, where a negative difference in Median Absolute Error and Quantile Loss at the 90<sup>th</sup> centile indicates better performance for cVAE, while positive differences in Spearman's correlation coefficient and Explained Variance indicate better performance for cVAE. Cliff's delta was used as the effect size measure. Permutation p-values reflect the proportion of permutations where the observed difference was as extreme as or more extreme than the actual data, providing a measure of statistical significance. Abbreviations: cVAE = conditional Variational Autoencoder; GAMLSS = Generalised Additive Models for Location, Scale and Shape; MFPR = Multivariate Fractional Polynomial Regression; HBR = Hierarchical Bayesian Regression.

In terms of Median Absolute Error, the differences between cVAE and other models were minor, with small effect sizes and high p-values. The most notable difference was between cVAE and MFPR with an observed median difference of −0.038; however, this difference did not reach statistically significance (p = 0.423). For the Quantile Loss at the 90th Percentile, cVAE showed a statistically significant improvement over MFPR (observed median difference = −0.046, effect size = −0.300, p-value = 0.025). Comparisons between cVAE and GAMLSS or HBR on this metric did not show statistically significant differences. Similarly, Spearman’s Correlation Coefficient and Explained Variance metrics indicated minimal and non-significant differences between cVAE and other models, reflecting comparable performance in capturing overall trends and variance. However, all models struggled to explain variance in the cerebellum, with values close to zero or even negative, as shown in Table S5. For instance, the explained variance of the left cerebellum was −0.01 for cVAE, 0.00 for GAMLSS and MFPR, and −0.07 for HBR. A similar pattern was observed in the left posterior artery callosal (PAC), with explained variance values of 0.05 for GAMLSS, 0.03 for both cVAE and MFPR, and −0.21.

Figure 3 (A-C) presents Bland-Altman plots that depict the distribution of differences in Median Absolute Error between cVAE and the other models across different brain regions. While some region-specific variations are apparent, the majority of brain regions cluster around the median difference lines (0.00, −0.01, and 0.01 for GAMLSS, MFPR, and HBR comparisons respectively), suggesting generally comparable performance. The distribution plot (Figure 3D) further supports this observation of comparable performance across models, with all four models showing similar central tendencies in their Median Absolute Error (cVAE: 0.52, GAMLSS: 0.53, MFPR: 0.56, HBR: 0.53). Additional Bland-Altman plots for other performance metrics are available in Figures S4–S6.

**Figure 3.**
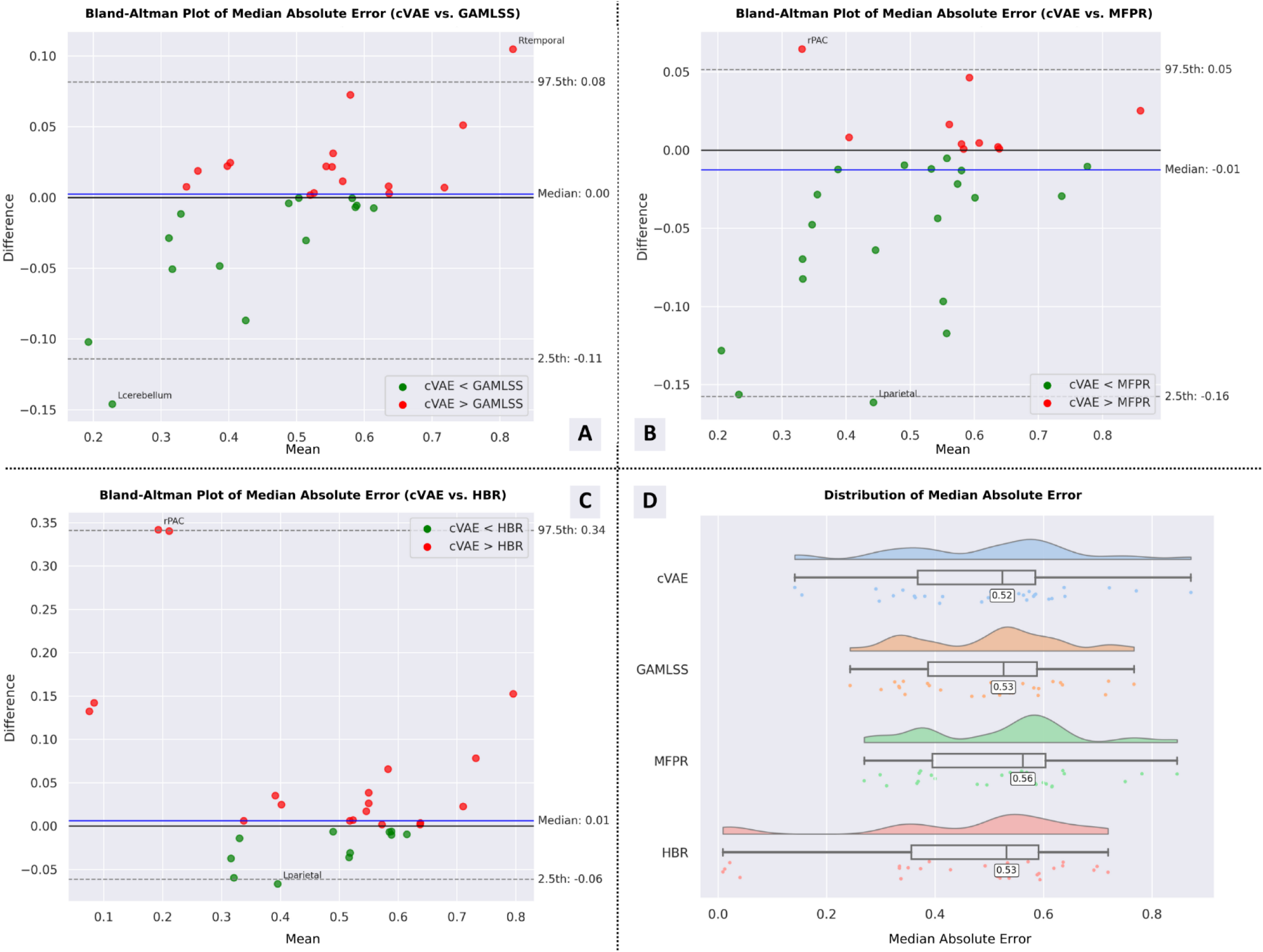
Bland-Altman Plots and Distribution of Median Absolute Error for cVAE vs Other Models on the Hold-Out Dataset. (A-C) Bland-Altman plots comparing the Median Absolute Error of the conditional Variational Autoencoder (cVAE) model against the Generalised Additive Models for Location, Scale, and Shape (GAMLSS) (A); Multivariate Fractional Polynomial Regression (MFPR) (B); and Hierarchical Bayesian Regression (HBR) (C) models. Each point represents a different brain region. Green points indicate regions where cVAE exhibits lower Median Absolute Error compared to the corresponding model, while red points show regions where the other model performs better. (D) Distribution of Median Absolute Error across all four models (cVAE, GAMLSS, MFPR, and HBR), visualising the variability and central tendency for each model’s performance. This metric is computed on the hold-out dataset.

### 3.4 Deviation Analysis

#### 3.4.1 Age Correlation Analysis on Hold-Out Dataset

Figure 4 illustrates the relationship between age and z-scores for whole-brain WMH volume on the hold-out dataset. As shown, Spearman correlation coefficients were consistently low across all models (cVAE: r = −0.02, GAMLSS: r = 0.03, MFPR: r = 0.09, HBR: r = 0.07), indicating effective control of age-related effects in whole-brain predictions. A more detailed summary of the correlations between age and z-scores across different brain regions is presented in Table S6. Most regions exhibited weak correlations, suggesting robust age-independent predictions. However, there were some notable exceptions, particularly in the cerebellum and posterior artery callosal (PAC) regions. In the cerebellum, cVAE showed relatively strong positive correlations (left: r = 0.473, right: r = 0.513), as did GAMLSS (left: r = 0.470, right: r = 0.125). The PAC regions displayed strong negative correlations for cVAE (left: r = −0.476, right: r = −0.547), while MFPR exhibited positive correlations (left: 0.305, right: 0.337).

**Figure 4.**
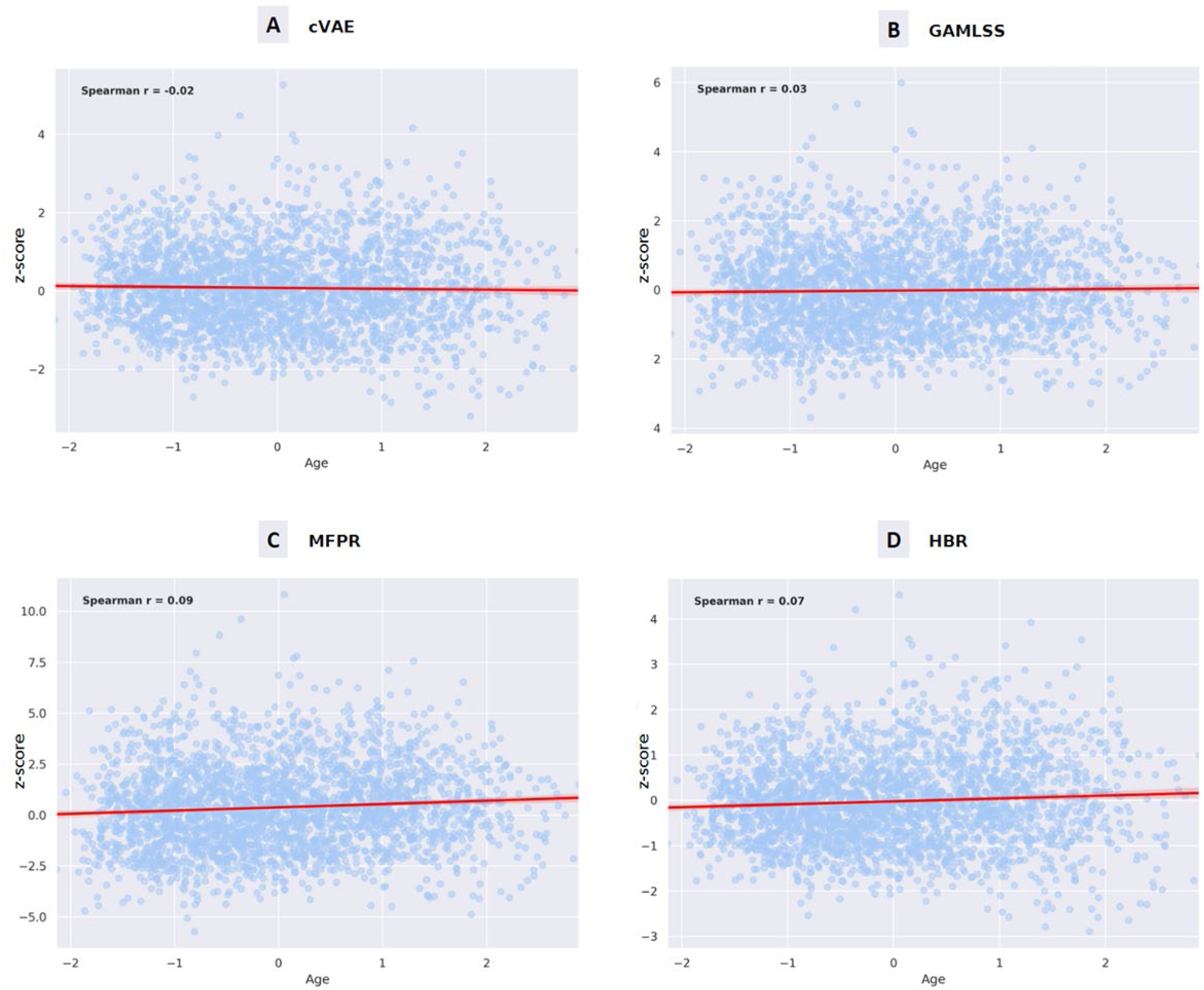
Correlation between Age and Z-Score of Whole-Brain White Matter Hyperintensity (WMH) Volume on the Hold-Out Dataset. This figure presents four scatter plots (A-D) showing the correlation between age and the residual whole-brain white matter hyperintensity (WMH) volume for different predictive models: cVAE, GAMLSS, MFPR, and HBR. Each plot displays the residual (observed minus predicted) WMH volume on the x-axis and age on the y-axis. Blue dots represent individual data points, while the red line indicates the trend with a shaded area showing the confidence interval. Low correlation between age and z-score indicates good control of age-related covariates in the model. Abbreviations: cVAE = conditional Variational Autoencoder; GAMLSS = Generalised Additive Models for Location, Scale and Shape; MFPR = Multivariate Fractional Polynomial Regression; HBR = Hierarchical Bayesian Regression.

#### 3.4.2 Extreme Deviation Analysis on Hold-Out Dataset

We examined the percentage of extreme deviations (z-score > 2.58) across brain regions and models using a hold-out dataset derived from the normotensive sample, as illustrated in Figure 5-6 and detailed in Table S7. As shown in Figure 5, MFPR consistently exhibited the highest frequency of extreme deviations across most regions. In contrast, cVAE and HBR demonstrated minimal deviations, often reporting 0% or negligible percentages. Figure 6 provides a visual representation of the spatial distribution of these deviations, further emphasising the trend in which MFPR displays greater deviations compared to cVAE and HBR across most brain regions.

**Figure 5.**
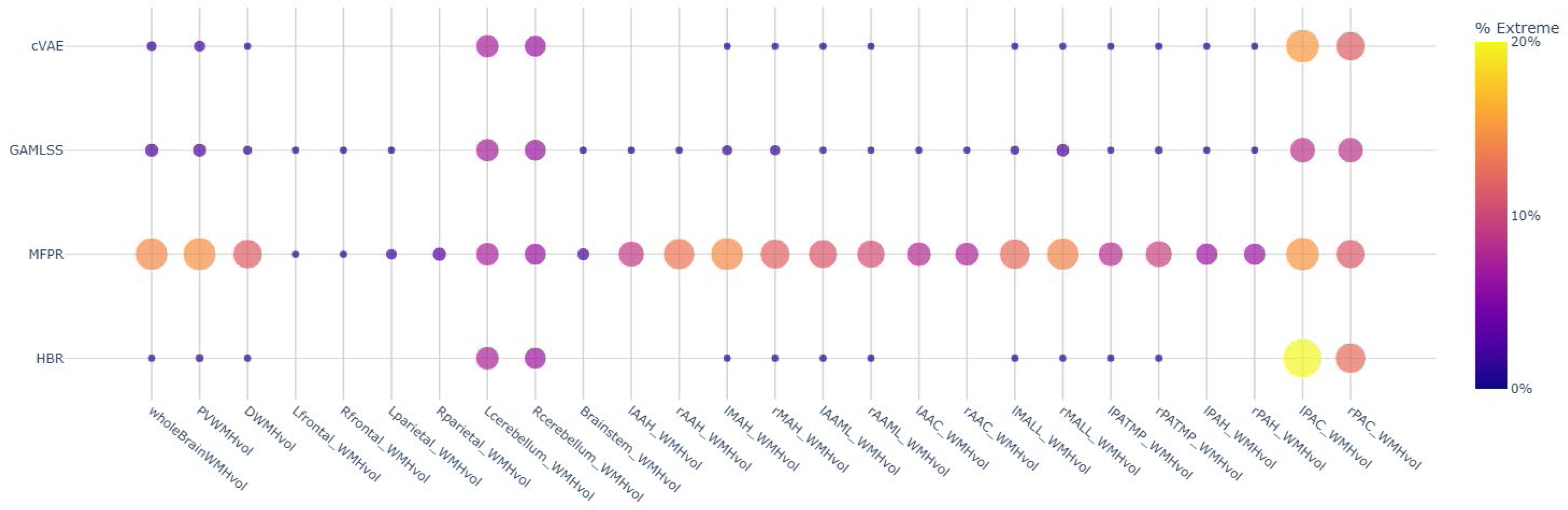
Percentage of Extreme Deviations on the Hold-Out Dataset across Models and Features. This bubble chart displays the percentage of samples with extreme deviations (z > 2.58) across models in a hold-out dataset. Bubble size and colour intensity indicate the percentage of extreme deviations, with larger, brighter bubbles representing higher percentages. Absent bubbles signify no extreme deviations. Low percentages of extreme deviations are desirable, as a good model should minimise extreme deviations on a hold-out dataset. Abbreviations: cVAE = conditional Variational Autoencoder; GAMLSS = Generalised Additive Models for Location, Scale and Shape; MFPR = Multivariate Fractional Polynomial Regression; HBR = Hierarchical Bayesian Regression; WMH = White Matter Hyperintensity. See Table S1 for abbreviations of lobar and arterial regions.

**Figure 6.**
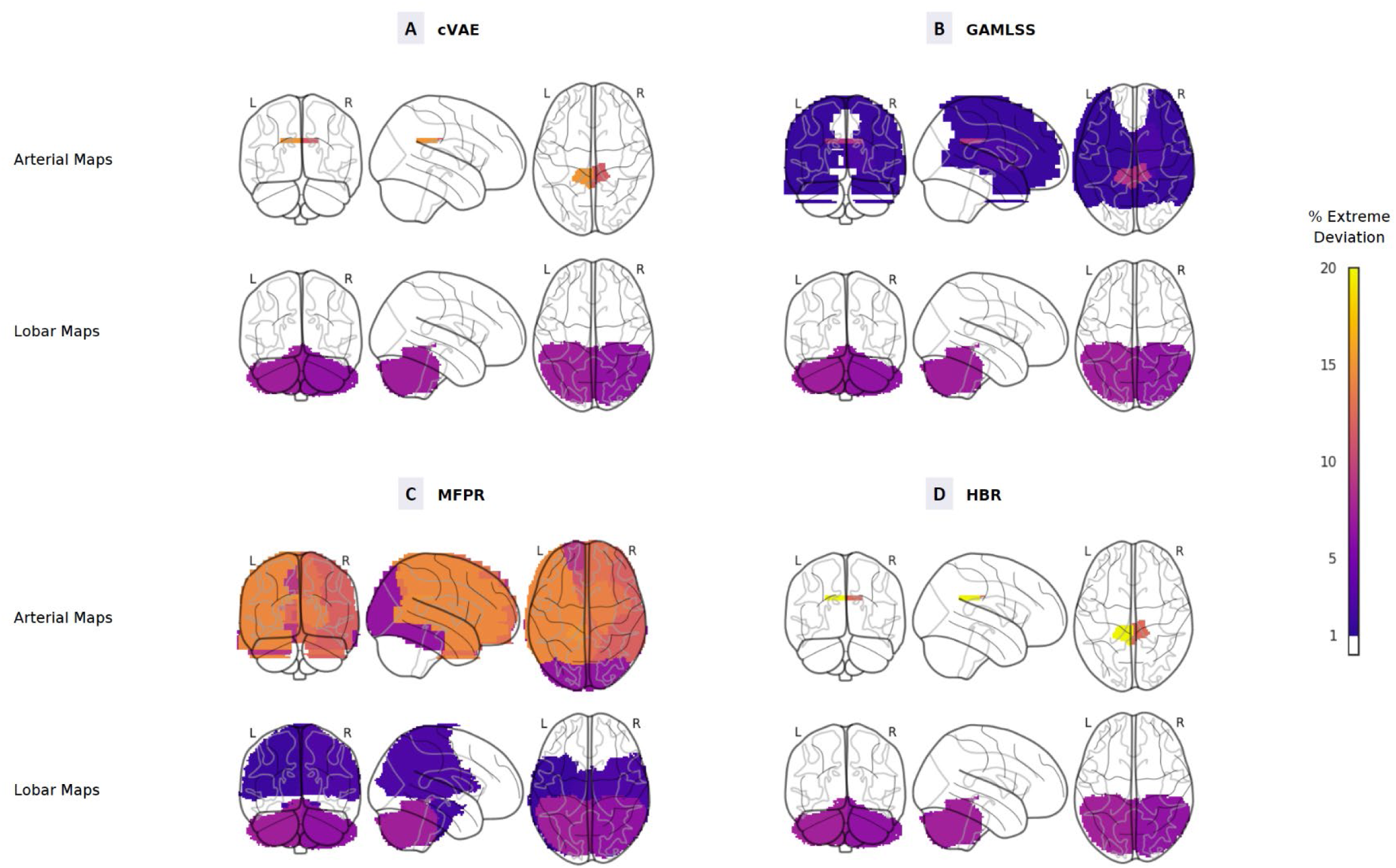
Spatial Distribution of Extreme Deviations on the Hold-Out Dataset across Models. This brain map displays the percentage of samples with extreme deviations (z > 2.58) across different models on the hold-out dataset, using arterial territory-specific and lobar parcellation. Extreme deviations indicate areas where model predictions diverge significantly from expected values, which is undesirable for accurate model generalisation. Colour intensity indicates the percentage of extreme deviations, with brighter colours representing higher percentages. The brain is shown in three views (coronal, sagittal, axial) for each model. Abbreviations: cVAE = conditional Variational Autoencoder; GAMLSS = Generalised Additive Models for Location, Scale and Shape; MFPR = Multivariate Fractional Polynomial Regression; HBR = Hierarchical Bayesian Regression. L and R denote Left and Right hemispheres respectively.

An exception to this pattern was observed in the PAC region, where HBR recorded the highest percentage of extreme deviations (left: 19.91%, right: 12.47%), followed by cVAE (left: 15.08%, right: 11.54%). Similarly, all models recorded relatively high percentages of extreme deviations in the cerebellum, ranging from 7.25% to 7.40% in the left cerebellum and 6.39% to 6.47% in the right cerebellum.

#### 3.4.2 Clinical Application

Figure 7 illustrates the distribution of whole-brain WMH volumes z-scores across hypertension levels (0-3) for all four models (cVAE, GAMLSS, MFPR, and HBR). Notably, all models displayed a similar trend: as hypertension levels increased, z-scores also rose. Details are shown in Figure S8 and Table S8, which presents Spearman’s correlations between z-scores and hypertension levels across brain regions. Most brain regions showed statistically significant positive correlations (p < 0.001), though the correlation coefficients were generally low, with values mostly below 0.1. For whole-brain WMH volume, correlation coefficients range from 0.077 (cVAE) to 0.093 (HBR), indicating a weak but consistent association between model-derived z-scores and hypertension severity.

**Figure 7.**
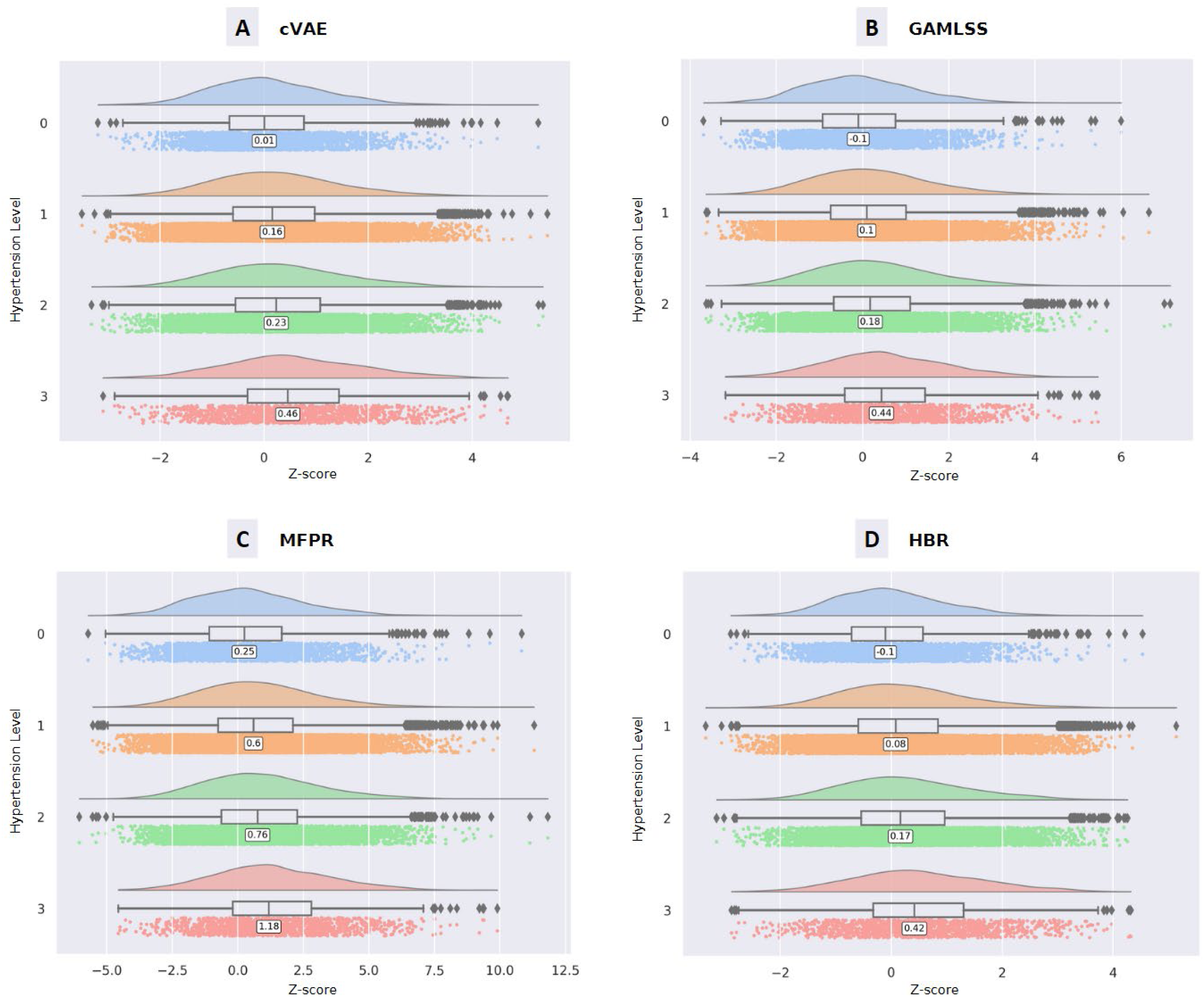
Comparison of Z-Scores for Whole-Brain WMH Volume across Hypertensive Levels and Models. (A-D) Box plots showing the distribution of Z-scores for whole-brain WMH volume stratified by hypertensive levels (0-3). Each panel corresponds to a different model for WMH volume estimation: (A) Conditional Variational Autoencoder (cVAE); (B) Generalised Additive Models for Location, Scale and Shape (GAMLSS); (C) Multivariate Fractional Polynomial Regression (MFPR); and (D) Hierarchical Bayesian Regression (HBR).

Certain regions, however, exhibited distinct patterns. For instance, the cerebellum showed weak and often non-significant correlations. In the left cerebellum, correlation values ranged from −0.015 (MFPR, p = 0.036) to 0.093 (cVAE, p < 0.001), with similar variability observed in the right cerebellum. In addition, the brain stem demonstrated relatively weak correlations, with coefficients ranging from 0 (GAMLSS, p = 0.970) to 0.030 (MFPR, p < 0.001). The PAC region presented an interesting case, with negative correlations for GAMLSS (left: −0.062, right: −0.074) and cVAE (left: −0.063, right: −0.091).

Figure 8 complements these findings by visualising the percentage of extreme deviations (z-score > 2.58) across hypertension levels for whole-brain, periventricular, and deep white matter WMH volumes. All models demonstrated an increase in extreme deviations with higher hypertension levels, particularly in whole-brain and periventricular WMH volumes. This trend was most pronounced in the MFPR model, which showed a significant increase in extreme deviations from hypertension level 0 to level 3. In contrast, the cVAE and HBR models displayed the most modest increase, while GAMLSS exhibited an intermediate pattern.

**Figure 8.**
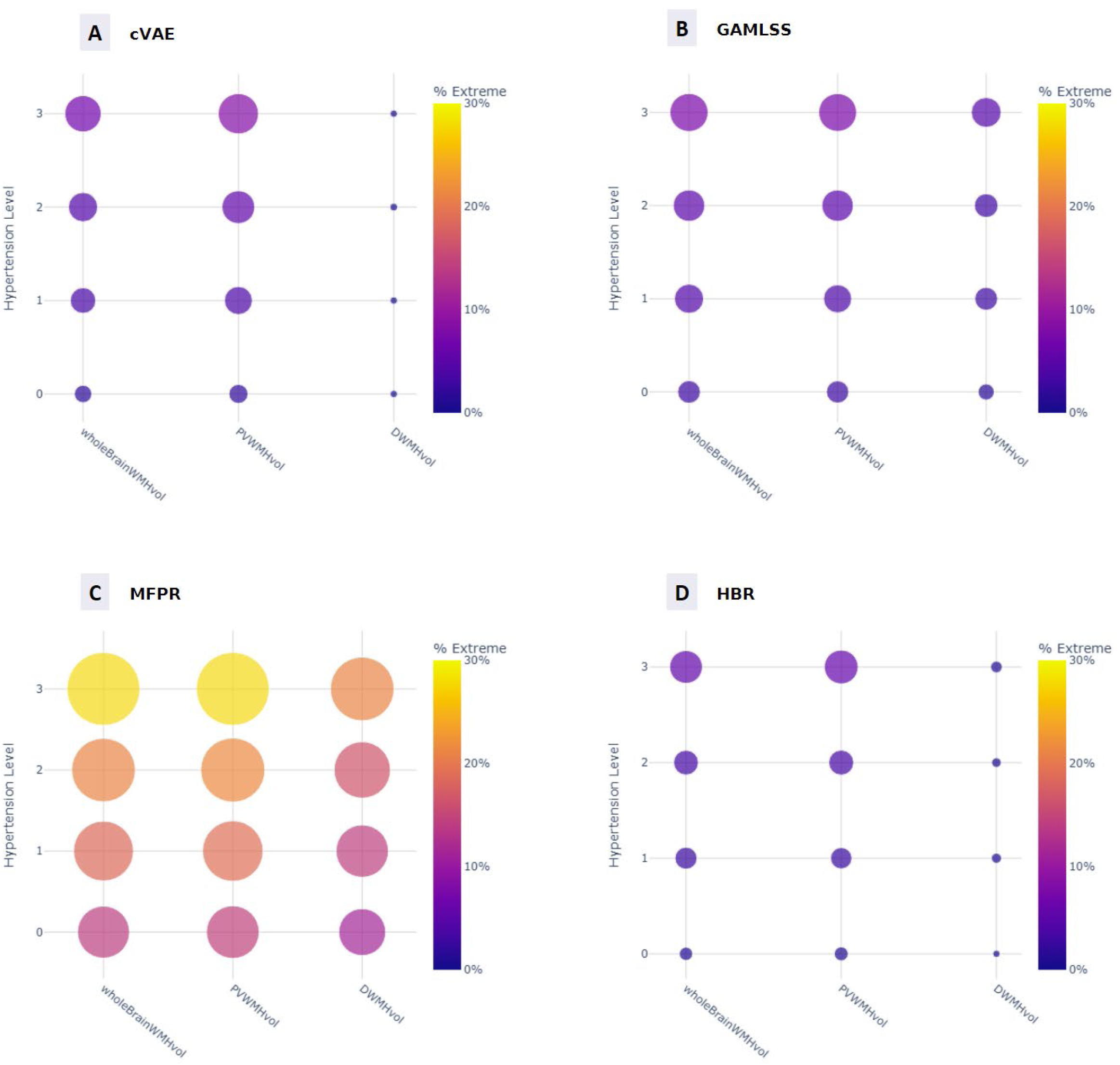
Comparison of Extreme Deviations across Models and Hypertension Levels. This bubble chart shows the percentage of samples with extreme deviations (z-score > 2.58) across hypertension levels (0-3) for 4 models: (A) Conditional Variational Autoencoder (cVAE); (B) Generalised Additive Models for Location, Scale and Shape (GAMLSS); (C) Multivariate Fractional Polynomial Regression (MFPR); and (D) Hierarchical Bayesian Regression (HBR). Bubble size and colour intensity indicate the percentage of extreme deviations, with larger, brighter bubbles representing higher percentages. Abbreviations: wholeBrainWMHvol = whole-brain WMH volume; PVWMHvol = periventricular WMH volume; DWMHvol = deep white matter hyperintensity volume.

Table S9 reveals further details, indicating that the cerebellum and PAC regions showed higher percentages of extreme deviations across all models and hypertension levels. In the cerebellum, the proportion of extreme deviations ranged from approximately 6% to 8.5% across all models and hypertension levels, with minimal variations between hypertension levels. The PAC regions displayed even higher percentages, ranging from about 7% to 35% depending on the model and hypertension level. Table S9 also showed that cVAE and HBR were more conservative in identifying extreme deviations, frequently showing 0% in regions where GAMLSS and MFPR detected some deviations. Conversely, MFPR consistently exhibited the highest percentages across most brain regions, with values reaching up to 27.89% in whole-brain analysis and over 20% in several subcortical areas.

### 3.5 Computational Efficiency

The computational efficiency of the models was assessed by comparing their elapsed times during the training and inference phases, as detailed in Table 4. It is worth noting that these times reflect different computational setups: cVAE and HBR utilised 48 CPU cores, with cVAE also employing 4 GPU for parallel processing, while GAMLSS and MFPR were run on standard configurations. Moreover, the Optuna framework was used for hyperparameter tuning of the cVAE model during the training process. The results reveal significant variations in computational requirements among the models. MFPR was the most efficient, with a total elapsed time of just 14 seconds, followed closely by GAMLSS at 57 seconds. In contrast, cVAE and HBR required substantially more computational resources. cVAE took 10,297 seconds (approximately 2.86 hours), while HBR demanded the most time at 15,161 seconds (about 4.21 hours).

**Table 4.**
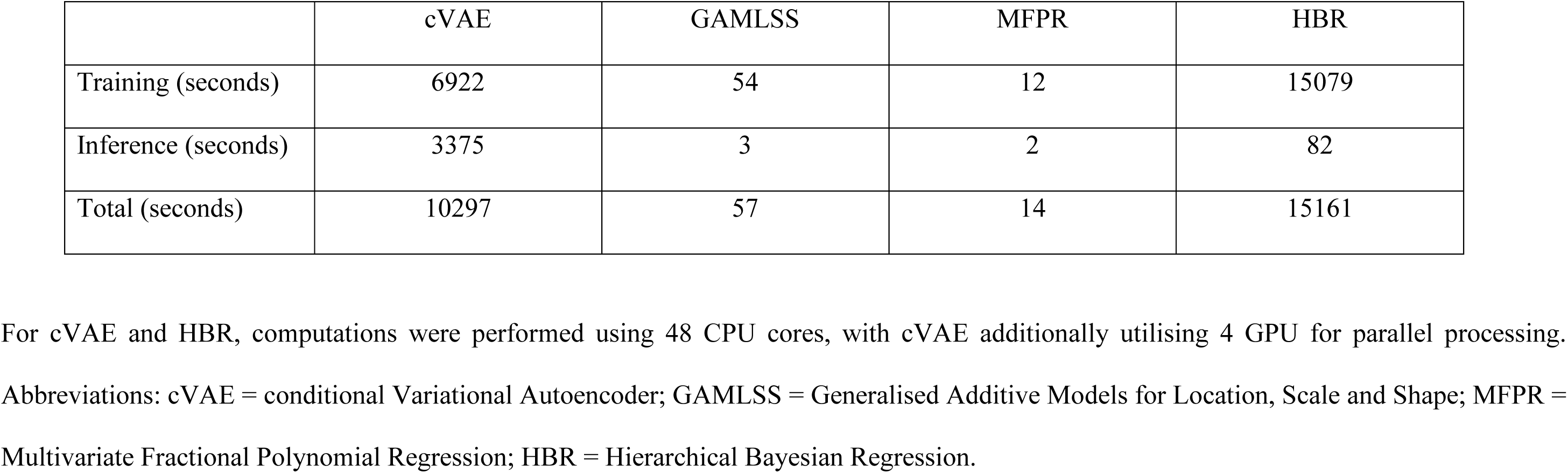
Elapsed time comparison across models during training and inference.

## 4. Discussion

A key contribution of this study is the development and refinement of the cVAE-based normative model, with a particular focus on integrating probabilistic inference. Our approach improves upon existing autoencoder-based approaches by enabling the model to directly predict distributions of neuroimaging data conditioned on covariates, rather than relying solely on reconstruction errors. This shift allows the model to produce more precise and nuanced probabilistic predictions, offering a deeper understanding of individual variations in brain structure. Importantly, this approach also addresses the limitations faced by traditional statistical normative models, which often struggle with handling high-dimensional data and capturing complex relationships. By leveraging the strengths of deep learning within a structured probabilistic framework, our model provides a more robust and flexible tool for analysing neuroimaging data.

Our results indicate that the proposed cVAE-based normative model performs comparably to well-established statistical methods (GAMLSS, MFPR and HBR) across all performance metrics. Notably, cVAE demonstrated a statistically significant improvement over MFPR in Quantile Loss at the 90^th^ percentile, reflecting its capacity to accurately model the upper range of deviations. Additionally, the z-score analysis on the hold-out dataset confirmed that our framework effectively produces age-independent deviations across most brain regions, successfully controlling for covariates and providing clinically relevant insights. The analysis of extreme deviations on the hold-out dataset further supports this, as most regions showed 0% extreme deviations, indicating that the cVAE excels in minimising false positives and delivering reliable predictions.

While cVAE and the other models generally performed well across most features, challenges were observed in certain brain regions, particularly in the cerebellum and posterior artery callosal (PAC) areas. Specifically, these regions had higher percentages of extreme deviations on the hold-out dataset across all models. These regions also exhibited higher correlations between z-scores and age, suggesting that age-related effects were not fully controlled. A likely explanation is that WMH volumes in the cerebellum and PAC regions are often zero or near-zero, leading to sparse data in the training set. This scarcity makes it difficult for the models to distinguish between true abnormalities and noise, resulting in higher rates of false positives. This limitation underscores the need for tailored approaches in normative models when dealing with sparse data.

Beyond challenges with sparse data, our analysis also highlighted a trade-off between model complexity and computational efficiency in the cVAE. Both the cVAE and HBR required several hours for training and inference, compared to mere minutes for GAMLSS or MFPR. However, the cVAE’s ability to handle complex, high-dimensional data compensates for this increased computational cost. Moreover, deep learning models like the cVAE can leverage GPUs for parallel processing, providing a pathway to reduce computational demands.

The clinical applicability of our approach was validated by the observed trends in WMH z-scores across varying levels of hypertension. All models, including the cVAE, consistently showed an increase in z-scores and the percentage of extreme deviations as hypertension severity increased. Although the correlations between WMH z-scores and hypertension levels were generally weak (with coefficients typically below 0.1), these associations were statistically significant across most brain regions. This indicates that cVAE can effectively capture clinically meaningful patterns. However, differences in model sensitivity should be noted. For example, the MFPR model identified a higher proportion of extreme deviations than the cVAE and other models, suggesting greater sensitivity to subtle abnormalities. However, this could also imply a higher rate of false positives, potentially leading to the over-identification of deviations without clinical significance. Conversely, the cVAE’s more conservative detection of extreme deviations may offer a better balance between sensitivity and specificity, providing a reliable method for identifying meaningful deviations without overestimating their prevalence.

Future work should focus on several key areas. First, there is a need to refine the model for regions with sparse data to reduce false positives. This may involve applying region-specific adjustments or incorporating additional predictors to improve accuracy. Second, to better handle higher-dimensional data, future iterations of the cVAE framework could be extended to a convolutional neural network (CNN) architecture. This would allow the model to analyse complex spatial structures, such as brain voxels, and capture more detailed patterns in neuroimaging data. Finally, future work should focus on enhancing the interpretability of cVAE models, potentially through explainable AI techniques, which could clarify the contributions of specific variables to the model’s predictions.

## 5. Conclusion

In conclusion, this study demonstrates that our proposed cVAE-based normative modelling framework offers a promising balance between the complexity of deep learning approaches and the practical needs of clinical neuroimaging analysis. While challenges remain in terms of computational efficiency, performance in data-sparse regions, and model interpretability, the cVAE shows significant strengths in handling high-dimensional data and accurately identifying clinically meaningful deviations. These attributes make the cVAE a strong candidate for advancing neuroimaging research, particularly in clinical applications where precision and reliability are critical.

## 6. Data and Code Availability

The UK Biobank dataset is available to researchers through a formal application process (https://www.ukbiobank.ac.uk/). The code of our enhanced cVAE framework is available at https://github.com/maiho24/BrainNormativeCVAE.git. Our implementation extends the normative modelling framework by (Lawry Aguila et al., 2022) (https://github.com/alawryaguila/normativecVAE), with refinements in both the model architecture and inference approach.

## 7. Acknowledgements

This study was conducted using resources and services provided by the National Computational Infrastructure (NCI Australia), which is supported by the Australian Government. We would also like to acknowledge the valuable assistance of Lei Fan in reviewing the model development section of this work.

