## Supplementary for "An Enhanced Conditional Variational Autoencoder-Based Normative Model for Neuroimaging Analysis"

### Supplementary Tables

#### Table S1 List of features and corresponding abbreviations.

| Feature | Definition |
|---|---|
| **General measures:** | |
| - wholeBrainWMHvol | Total volume of WMH in the entire brain ($mm^3$) |
| - PVWMHvol | WMH volume of the periventricular areas ($mm^3$) |
| - DWMHvol | Total volume of deep WMH ($mm^3$) |
| **Lobe-specific measures:** | |
| - lFrontal_WMHvol | WMH volume of the left frontal lobe ($mm^3$) |
| - rFrontal_WMHvol | WMH volume of the right frontal lobe ($mm^3$) |
| - lTemporal_WMHvol | WMH volume of the left temporal lobe ($mm^3$) |
| - rTemporal_WMHvol | WMH volume of the right temporal lobe ($mm^3$) |
| - lParietal_WMHvol | WMH volume of the left parietal lobe ($mm^3$) |
| - rParietal_WMHvol | WMH volume of the right parietal lobe ($mm^3$) |
| - lOccipital_WMHvol | WMH volume of the left occipital lobe ($mm^3$) |
| - rOccipital_WMHvol | WMH volume of the right occipital lobe ($mm^3$) |
| - lCerebellum_WMHvol | WMH volume of the left cerebellum ($mm^3$) |
| - rCerebellum_WMHvol | WMH volume of the right cerebellum ($mm^3$) |
| - Brainstem_WMHvol | WMH volume of the brainstem ($mm^3$) |

| | |
|---|---|
| **Arterial territory-specific measures:** | |
| - lAAH_WMHvol | WMH volume of the left anterior artery hemisphere (mm$^3$) |
| - rAAH_WMHvol | WMH volume of the right anterior artery hemisphere (mm$^3$) |
| - lMAH_WMHvol | WMH volume of the left middle artery hemisphere (mm$^3$) |
| - rMAH_WMHvol | WMH volume of the right middle artery hemisphere (mm$^3$) |
| - lAAML_WMHvol | WMH volume of the left anterior medial lenticulostriate (mm$^3$) |
| - rAAML_WMHvol | WMH volume of the right anterior medial lenticulostriate (mm$^3$) |
| - lAAC_WMHvol | WMH volume of the left anterior callosal (mm$^3$) |
| - rAAC_WMHvol | WMH volume of the right anterior callosal (mm$^3$) |
| - lMALL_WMHvol | WMH volume of the left middle artery lateral lenticulostriate (mm$^3$) |
| - rMALL_WMHvol | WMH volume of the right middle artery lateral lenticulostriate (mm$^3$) |
| - lPATMP_WMHvol | WMH volume of the left posterior artery thalamic and midbrain perforators (mm$^3$) |
| - rPATMP_WMHvol | WMH volume of the right posterior artery thalamic and midbrain perforators (mm$^3$) |
| - lPAH_WMHvol | WMH in the left posterior artery hemisphere (mm$^3$) |
| - rPAH_WMHvol | WMH in the right posterior artery hemisphere (mm$^3$) |
| - lPAC_WMHvol | WMH in the left posterior artery callosal (mm$^3$) |
| - rPAC_WMHvol | WMH in the left posterior artery callosal (mm$^3$) |

Abbreviations: WMH = white matter hyperintensity.

**Table S2 List of exclusion codes and corresponding diseases from UK Biobank dataset.**

| **Code** | **Disease** |
|---|---|
| 1081 | stroke |
| 1082 | transient ischaemic attack (tia) |
| 1083 | subdural haemorrhage/haematoma |
| 1086 | subarachnoid haemorrhage |
| 1240 | neurological injury/trauma |
| 1243 | psychological/psychiatric problem |
| 1244 | infection of nervous system |
| 1245 | brain abscess/intracranial abscess |
| 1246 | encephalitis |
| 1247 | meningitis |
| 1256 | acute infective polyneuritis/guillain-barre syndrome |
| 1258 | chronic/degenerative neurological problem |
| 1259 | motor neurone disease |
| 1261 | multiple sclerosis |
| 1262 | Parkinsons disease |
| 1263 | dementia/Alzheimer's/cognitive impairment |
| 1264 | epilepsy |
| 1266 | head injury |
| 1289 | schizophrenia |
| 1291 | mania/bipolar disorder/manic depression |

| | |
|---|---|
| 1297 | muscle/soft tissue problem |
| 1425 | cerebral aneurysm |
| 1433 | cerebral palsy |
| 1491 | brain haemorrhage |
| 1524 | spina bifida |
| 1583 | ischaemic stroke |
| 1659 | meningioma / benign meningeal tumour |

**Table S3 Descriptive statistics of WMH volumes (in mm$^3$) from UBO Detector in the training and evaluation sample.**

| Feature | Training sample | Evaluation sample | | | |
|---|---|---|---|---|---|
| | All training mean ± SD | Normotensive mean ± SD | Hypertensive Level 1 mean ± SD | Hypertensive Level 2 mean ± SD | Hypertensive Level 3 mean ± SD |
| **General measures:** | | | | | |
| - wholeBrainWMHvol | 1761.92 ±2474.39 | 1761.5 ±3241.66 | 2433.40 ±3315.97 | 2826.63 ±3763.55 | 3933.86 ±5122.58 |
| - PVWMHvol | 1304.62 ±1686.83 | 1294.22 ±2069.13 | 1797.52 ±2373.29 | 2099.64 ±2755.11 | 2918.03 ±3608.72 |
| - DWMHvol | 434.93 ±958.26 | 442.23 ±1245.05 | 608.44 ±1140.08 | 697.37 ±1226.06 | 980.13 ±1929.39 |
| **Lobe-specific measures:** | | | | | |
| - lFrontal_WMHvol | 86.09 ±238.28 | 86.38 ±226.60 | 120.00 ±263.27 | 132.58 ±269.40 | 174.69 ±351.04 |
| - rFrontal_WMHvol | 91.27 ±228.82 | 90.27 ±246.64 | 127.78 ±298.69 | 141.77 ±270.57 | 187.73 ±369.43 |
| - lTemporal_WMHvol | 12.09 ±36.09 | 13.89 ±98.36 | 17.17 ±54.60 | 19.52 ±51.12 | 28.83 ±80.48 |
| - rTemporal_WMHvol | 10.50 ±28.30 | 14.45 ±159.45 | 14.68 ±43.10 | 17.96 ±82.08 | 25.05 ±62.13 |
| - lParietal_WMHvol | 82.2 ±260.33 | 83.2 ±291.76 | 121.44 ±309.61 | 142.83 ±376.07 | 216.93 ±577.52 |
| - rParietal_WMHvol | 84.63 ±250.14 | 84.04 ±322.32 | 124.34 ±302.58 | 148.21 ±331.60 | 232.31 ±628.92 |
| - lOccipital_WMHvol | 26.90 ±44.59 | 28.59 ±48.70 | 33.48 ±55.43 | 38.33 ±64.68 | 46.90 ±75.79 |
| - rOccipital_WMHvol | 41.19 ±62.23 | 41.41 ±59.57 | 49.53 ±71.76 | 56.16 ±81.59 | 67.68 ±99.88 |
| - lCerebellum_WMHvol | 0.36 ±5.85 | 0.33 ±2.75 | 0.48 ±10.46 | 0.46 ±4.52 | 0.42 ±4.50 |
| - rCerebellum_WMHvol | 0.27 ±3.63 | 0.33 ±3.35 | 0.35 ±3.30 | 0.41 ±6.04 | 0.35 ±3.42 |
| - Brainstem_WMHvol | 10.22 ±20.04 | 12.13 ±81.28 | 13.26 ±33.23 | 14.59 ±36.23 | 17.50 ±46.67 |
| **Arterial territory-specific measures:** | | | | | |
| - lAAH_WMHvol | 55.05 ±110.58 | 56.09 ±160.71 | 76.02 ±133.11 | 85.49 ±141.58 | 119.27 ±196.62 |
| - rAAH_WMHvol | 50.89 ±110.19 | 53.3 ±186.67 | 72.51 ±161.27 | 83.76 ±167.70 | 110.29 ±194.57 |

| | | | | | | |
|---|---|---|---|---|---|---|
| - | lMAH_WMHvol | 245.22 ±623.30 | 247.84 ±731.59 | 368.39 ±794.88 | 429.90 ±888.72 | 627.57 ±1244.33 |
| - | rMAH_WMHvol | 275.85 ±589.42 | 271.33 ±757.22 | 389.41 ±710.57 | 457.38 ±803.76 | 655.03 ±1126.29 |
| - | lAAML_WMHvol | 92.79 ±100.80 | 91.46 ±110.80 | 117.37 ±133.18 | 135.59 ±152.80 | 177.49 ±196.81 |
| - | rAAML_WMHvol | 96.09 ±101.38 | 92.64 ±108.94 | 119.87 ±131.67 | 136.80 ±152.67 | 176.91 ±191.24 |
| - | lAAC_WMHvol | 103.44 ±129.24 | 100.53 ±131.26 | 135.53 ±170.67 | 157.24 ±191.27 | 210.08 ±233.72 |
| - | rAAC_WMHvol | 92.43 ±112.32 | 91.49 ±113.95 | 117.62 ±145.95 | 131.62 ±161.35 | 170.03 ±190.67 |
| - | lMALL_WMHvol | 250.13 ±349.19 | 252.28 ±412.97 | 363.94 ±500.55 | 429.11 ±570.62 | 599.49 ±711.94 |
| - | rMALL_WMHvol | 221.45 ±316.64 | 218.71 ±350.43 | 321.28 ±446.68 | 376.84 ±507.64 | 526.30 ±632.23 |
| - | lPATMP_WMHvol | 60.11 ±37.45 | 60.28 ±68.44 | 59.75 ±39.53 | 61.23 ±42.60 | 64.20 ±48.73 |
| - | rPATMP_WMHvol | 42.42 ±29.66 | 42.52 ±61.31 | 43.82 ±33.49 | 45.52 ±37.60 | 55.66 ±181.31 |
| - | lPAH_WMHvol | 66.39 ±138.87 | 70.2 ±184.75 | 93.41 ±195.78 | 111.97 ±233.16 | 169.13 ±361.95 |
| - | rPAH_WMHvol | 93.58 ±195.84 | 94.49 ±236.67 | 130.50 ±256.44 | 156.03 ±294.98 | 230.98 ±476.39 |
| - | lPAC_WMHvol | 3.31 ±18.69 | 3.79 ±22.73 | 6.45 ±31.61 | 8.41 ±37.31 | 15.56 ±52.39 |
| - | rPAC_WMHvol | 1.40 ±10.50 | 1.29 ±10.95 | 2.70 ±16.86 | 3.39 ±19.54 | 6.41 ±27.78 |

Abbreviations: WMH = white matter hyperintensity; SD = standard deviation. See Table S1 for abbreviations of lobar and arterial regions.

**Table S4 Correlations between covariates and WMH volumes across brain regions using Spearman and point-biserial tests in the training dataset.**

| | Spearman Correlation Test | | | | Point-Biserial Correlation Test | | | | | | | | | |
|---|---|---|---|---|---|---|---|---|---|---|---|---|---|---|
| | Age | | Intracranial Volume | | Sex | | Diabetes | | Hypercholesterolemia | | Obesity | | Smoking | |
| | $r_s$ | *p*-value | $r_s$ | *p*-value | $r_{pb}$ | *p*-value | $r_{pb}$ | *p*-value | $r_{pb}$ | *p*-value | $r_{pb}$ | *p*-value | $r_{pb}$ | *p*-value |
| whole-brain | 0.398 | **<0.001** | 0.067 | **<0.001** | -0.021 | 0.119 | 0.055 | **<0.001** | 0.128 | **<0.001** | 0.016 | 0.467 | 0.071 | **<0.001** |
| periventricular | 0.374 | **<0.001** | 0.059 | **<0.001** | -0.044 | **0.001** | 0.051 | **<0.001** | 0.119 | **<0.001** | 0.015 | 0.486 | 0.073 | **<0.001** |
| deep white matter | 0.355 | **<0.001** | 0.065 | **<0.001** | 0.042 | **0.002** | 0.053 | **<0.001** | 0.112 | **<0.001** | 0.006 | 0.844 | 0.039 | **0.006** |
| lFrontal | 0.285 | **<0.001** | 0.039 | **0.004** | 0.040 | **0.003** | 0.053 | **<0.001** | 0.058 | **<0.001** | 0.002 | 0.952 | 0.046 | **0.001** |
| rFrontal | 0.278 | **<0.001** | 0.047 | **<0.001** | 0.036 | **0.009** | 0.055 | **<0.001** | 0.066 | **<0.001** | 0.009 | 0.661 | 0.024 | 0.074 |
| lTemporal | 0.201 | **<0.001** | 0.089 | **<0.001** | 0.108 | **<0.001** | 0.032 | **0.021** | 0.073 | **<0.001** | 0.017 | 0.467 | 0.032 | **0.026** |
| rTemporal | 0.201 | **<0.001** | 0.066 | **<0.001** | 0.109 | **<0.001** | 0.018 | 0.209 | 0.060 | **<0.001** | 0.021 | 0.407 | 0.030 | **0.040** |
| lParietal | 0.337 | **<0.001** | 0.018 | 0.281 | 0.027 | **0.048** | 0.043 | **0.003** | 0.081 | **<0.001** | 0.005 | 0.894 | 0.041 | **0.005** |
| rParietal | 0.346 | **<0.001** | 0.018 | 0.281 | 0.029 | **0.034** | 0.036 | **0.012** | 0.071 | **<0.001** | 0.016 | 0.467 | 0.020 | 0.139 |
| lOccipital | 0.132 | **<0.001** | 0.048 | **<0.001** | 0.058 | **<0.001** | 0.011 | 0.428 | 0.037 | **0.005** | -0.012 | 0.605 | 0.011 | 0.388 |
| rOccipital | 0.155 | **<0.001** | 0.080 | **<0.001** | 0.066 | **<0.001** | 0.008 | 0.569 | 0.029 | **0.034** | -0.011 | 0.605 | 0.021 | 0.121 |
| lCerebellum | -0.033 | **0.012** | 0.037 | **0.006** | 0.039 | 0.004 | 0.029 | **0.036** | 0.002 | 0.878 | 0.023 | 0.393 | 0.025 | 0.071 |
| rCerebellum | -0.020 | 0.120 | 0.046 | **<0.001** | 0.072 | **<0.001** | 0.013 | 0.356 | -0.004 | 0.789 | -0.019 | 0.407 | -0.020 | 0.142 |
| Brainstem | 0.124 | **<0.001** | 0.048 | **<0.001** | 0.097 | **<0.001** | 0.019 | 0.196 | 0.022 | 0.102 | 0.025 | 0.316 | 0.027 | 0.054 |
| lAAH | 0.312 | **<0.001** | 0.028 | **0.033** | 0.004 | 0.801 | 0.036 | **0.012** | 0.049 | **<0.001** | -0.001 | 0.952 | 0.029 | **0.042** |
| rAAH | 0.304 | **<0.001** | 0.024 | 0.065 | -0.002 | 0.885 | 0.030 | **0.035** | 0.049 | **<0.001** | 0.018 | 0.465 | 0.026 | 0.058 |

| | | | | | | | | | | | | | | |
|---|---|---|---|---|---|---|---|---|---|---|---|---|---|---|
| lMAH | 0.336 | **<0.001** | 0.051 | **<0.001** | 0.003 | 0.840 | 0.048 | **0.001** | 0.103 | **<0.001** | 0.001 | 0.952 | 0.048 | <0.001 |
| rMAH | 0.362 | **<0.001** | 0.048 | **<0.001** | -0.031 | **0.023** | 0.042 | **0.003** | 0.106 | **<0.001** | 0.025 | 0.316 | 0.050 | **<0.001** |
| lAAML | 0.351 | **<0.001** | -0.014 | 0.281 | -0.115 | **<0.001** | 0.047 | **0.001** | 0.085 | **<0.001** | 0.013 | 0.565 | 0.051 | **<0.001** |
| rAAML | 0.381 | **<0.001** | 0.03 | **0.025** | -0.087 | **<0.001** | 0.057 | **<0.001** | 0.094 | **<0.001** | 0.001 | 0.952 | 0.054 | **<0.001** |
| lAAC | 0.248 | **<0.001** | -0.032 | **0.016** | -0.117 | **<0.001** | 0.023 | 0.108 | 0.053 | **<0.001** | 0.015 | 0.486 | 0.040 | **0.006** |
| rAAC | 0.189 | **<0.001** | -0.047 | **<0.001** | -0.105 | **<0.001** | 0.014 | 0.329 | 0.009 | 0.533 | 0.011 | 0.622 | 0.031 | **0.033** |
| lMALL | 0.350 | **<0.001** | 0.071 | **<0.001** | -0.03 | **0.028** | 0.048 | **0.001** | 0.108 | **<0.001** | 0.02 | 0.407 | 0.079 | **<0.001** |
| rMALL | 0.306 | **<0.001** | 0.084 | **<0.001** | 0.017 | 0.221 | 0.041 | **0.003** | 0.107 | **<0.001** | 0.025 | 0.316 | 0.066 | **<0.001** |
| lPATMP | -0.017 | 0.191 | 0.067 | **<0.001** | -0.079 | **<0.001** | 0.003 | 0.826 | 0.011 | 0.441 | -0.006 | 0.844 | 0.029 | **0.042** |
| rPATMP | 0.038 | **0.003** | 0.075 | **<0.001** | -0.081 | **<0.001** | 0.019 | 0.190 | 0.021 | 0.115 | 0.002 | 0.952 | 0.026 | 0.064 |
| lPAH | 0.241 | **<0.001** | 0.064 | **<0.001** | 0.037 | **0.007** | 0.023 | 0.108 | 0.052 | **<0.001** | -0.032 | 0.316 | 0.016 | 0.235 |
| rPAH | 0.260 | **<0.001** | 0.090 | **<0.001** | 0.059 | **<0.001** | 0.007 | 0.636 | 0.056 | **<0.001** | -0.02 | 0.407 | 0.018 | 0.178 |
| lPAC | 0.187 | **<0.001** | 0.049 | **<0.001** | 0.012 | 0.397 | 0.033 | **0.021** | 0.072 | **<0.001** | 0.027 | 0.316 | 0.036 | **0.012** |
| rPAC | 0.138 | **<0.001** | 0.053 | **<0.001** | 0.023 | 0.086 | 0.033 | **0.021** | 0.073 | **<0.001** | 0.004 | 0.894 | 0.029 | **0.042** |

This table displays Spearman correlation coefficients ($r_s$) and Point-Biserial correlation coefficients ($r_{pb}$) and corresponding $p$-values for associations between various covariates and white matter hyperintensity (WMH) volumes across different brain regions in the training dataset. The Spearman correlation is used for assessing relationships between continuous variables, while the point-biserial correlation is used to assess relationships between binary variables and WMH volumes. Statistically significant correlations ($p < 0.05$) are highlighted. Abbreviations: GAMLSS = Generalised Additive Models for Location, Scale and Shape; MFPR = Multivariate Fractional Polynomial Regression; cVAE = conditional Variational Autoencoder; HBR = Hierarchical Bayesian Regression. See Table S1 for abbreviations of lobar and arterial regions.

**Table S5 Summary of model performance metrics across brain regions and performance metrics.**

| | Median Absolute Error | | | | Quantile Loss $90^{th}$ | | | | Spearman's Correlation | | | | Explained Variance | | | |
|---|---|---|---|---|---|---|---|---|---|---|---|---|---|---|---|---|
| | cVAE | GAM-LSS | MFPR | HBR | cVAE | GAM-LSS | MFPR | HBR | cVAE | GAM-LSS | MFPR | HBR | cVAE | GAM-LSS | MFPR | HBR |
| whole-brain | 0.58 | 0.58 | 0.58 | 0.59 | 0.37 | 0.34 | 0.42 | 0.34 | 0.45** | 0.46** | 0.45** | 0.46** | 0.21 | 0.22 | 0.18 | 0.22 |
| periventricular | 0.58 | 0.59 | 0.58 | 0.59 | 0.37 | 0.35 | 0.42 | 0.35 | 0.43** | 0.44** | 0.43** | 0.44** | 0.21 | 0.21 | 0.17 | 0.21 |
| deep white matter | 0.53 | 0.52 | 0.54 | 0.52 | 0.35 | 0.32 | 0.39 | 0.32 | 0.40** | 0.40** | 0.40** | 0.41** | 0.13 | 0.14 | 0.12 | 0.14 |
| lFrontal | 0.50 | 0.53 | 0.62 | 0.53 | 0.34 | 0.36 | 0.42 | 0.37 | 0.33** | 0.33** | 0.33** | 0.29** | 0.07 | 0.07 | 0.07 | 0.03 |
| rFrontal | 0.50 | 0.5 | 0.6 | 0.53 | 0.36 | 0.35 | 0.42 | 0.37 | 0.31** | 0.31** | 0.31** | 0.29** | 0.07 | 0.07 | 0.06 | 0.03 |
| lTemporal | 0.77 | 0.72 | 0.78 | 0.69 | 0.46 | 0.42 | 0.46 | 0.43 | 0.25** | 0.25** | 0.25** | 0.18** | 0.04 | 0.04 | 0.04 | 0.01 |
| rTemporal | 0.87 | 0.77 | 0.85 | 0.72 | 0.50 | 0.43 | 0.48 | 0.43 | 0.23** | 0.23** | 0.21** | 0.19** | 0.03 | 0.04 | 0.03 | 0.02 |
| lParietal | 0.36 | 0.41 | 0.52 | 0.43 | 0.28 | 0.32 | 0.39 | 0.32 | 0.38** | 0.39** | 0.39** | 0.39** | 0.07 | 0.10 | 0.10 | 0.05 |
| rParietal | 0.41 | 0.39 | 0.48 | 0.39 | 0.33 | 0.31 | 0.38 | 0.32 | 0.38** | 0.38** | 0.38** | 0.37** | 0.07 | 0.08 | 0.07 | 0.05 |
| lOccipital | 0.72 | 0.71 | 0.75 | 0.70 | 0.44 | 0.43 | 0.45 | 0.42 | 0.15** | 0.14** | 0.14** | 0.14** | 0.01 | 0.01 | 0.01 | 0.01 |
| rOccipital | 0.62 | 0.54 | 0.57 | 0.55 | 0.43 | 0.38 | 0.39 | 0.38 | 0.12** | 0.14** | 0.12** | 0.1** | 0 | 0.01 | 0.01 | 0 |
| lCerebellum | 0.15 | 0.30 | 0.31 | 0.01 | 0.25 | 0.25 | 0.25 | 0.24 | 0.03 | 0.03 | 0.02 | 0 | -0.01 | 0 | 0 | -0.07 |
| rCerebellum | 0.14 | 0.24 | 0.27 | 0.01 | 0.24 | 0.25 | 0.25 | 0.24 | 0.03 | 0.05* | 0.04* | 0 | -0.01 | 0 | 0 | -0.06 |
| Brainstem | 0.41 | 0.39 | 0.40 | 0.37 | 0.33 | 0.31 | 0.32 | 0.30 | 0.17** | 0.16** | 0.15** | 0.13** | 0.02 | 0.02 | 0.02 | 0.01 |
| lAAH | 0.30 | 0.33 | 0.37 | 0.33 | 0.22 | 0.26 | 0.31 | 0.27 | 0.36** | 0.35** | 0.35** | 0.34** | 0.04 | 0.05 | 0.04 | 0.04 |
| rAAH | 0.34 | 0.33 | 0.37 | 0.33 | 0.27 | 0.26 | 0.30 | 0.26 | 0.33** | 0.34** | 0.34** | 0.34** | 0.05 | 0.05 | 0.05 | 0.05 |
| lMAH | 0.49 | 0.49 | 0.50 | 0.49 | 0.33 | 0.31 | 0.36 | 0.30 | 0.37** | 0.37** | 0.36** | 0.37** | 0.10 | 0.11 | 0.09 | 0.11 |

| | | | | | | | | | | | | | | | | |
|---|---|---|---|---|---|---|---|---|---|---|---|---|---|---|---|---|
| rMAH | 0.55 | 0.53 | 0.56 | 0.54 | 0.39 | 0.33 | 0.40 | 0.34 | 0.4** | 0.4** | 0.4** | 0.4** | 0.11 | 0.12 | 0.1 | 0.12 |
| lAAML | 0.56 | 0.54 | 0.58 | 0.54 | 0.36 | 0.33 | 0.40 | 0.33 | 0.41** | 0.42** | 0.4** | 0.41** | 0.11 | 0.11 | 0.09 | 0.11 |
| rAAML | 0.52 | 0.52 | 0.56 | 0.51 | 0.33 | 0.32 | 0.40 | 0.32 | 0.45** | 0.46** | 0.45** | 0.45** | 0.19 | 0.20 | 0.16 | 0.20 |
| lAAC | 0.57 | 0.56 | 0.59 | 0.57 | 0.32 | 0.32 | 0.37 | 0.33 | 0.31** | 0.31** | 0.29** | 0.31** | 0.08 | 0.09 | 0.07 | 0.09 |
| rAAC | 0.57 | 0.54 | 0.55 | 0.53 | 0.38 | 0.32 | 0.35 | 0.32 | 0.24** | 0.25** | 0.24** | 0.24** | 0.02 | 0.05 | 0.04 | 0.04 |
| lMALL | 0.59 | 0.59 | 0.62 | 0.59 | 0.37 | 0.35 | 0.42 | 0.35 | 0.43** | 0.44** | 0.43** | 0.43** | 0.19 | 0.20 | 0.15 | 0.19 |
| rMALL | 0.61 | 0.62 | 0.61 | 0.62 | 0.43 | 0.37 | 0.43 | 0.37 | 0.35** | 0.36** | 0.35** | 0.36** | 0.12 | 0.15 | 0.11 | 0.15 |
| lPATMP | 0.64 | 0.63 | 0.64 | 0.64 | 0.35 | 0.39 | 0.40 | 0.39 | 0.12** | 0.14** | 0.13** | 0.15** | 0 | 0.01 | 0.01 | 0.02 |
| rPATMP | 0.64 | 0.63 | 0.64 | 0.64 | 0.36 | 0.40 | 0.41 | 0.40 | 0.15** | 0.17** | 0.17** | 0.16** | 0.01 | 0.02 | 0.02 | 0.02 |
| lPAH | 0.29 | 0.34 | 0.37 | 0.35 | 0.19 | 0.26 | 0.29 | 0.27 | 0.27** | 0.26** | 0.25** | 0.26** | -0.01 | 0.03 | 0.03 | 0.02 |
| rPAH | 0.32 | 0.33 | 0.37 | 0.34 | 0.23 | 0.25 | 0.29 | 0.25 | 0.27** | 0.27** | 0.26** | 0.26** | 0.03 | 0.03 | 0.02 | 0.03 |
| lPAC | 0.38 | 0.47 | 0.39 | 0.04 | 0.41 | 0.39 | 0.40 | 0.45 | 0.22** | 0.21** | 0.19** | -0.04* | 0.03 | 0.05 | 0.03 | -0.21 |
| rPAC | 0.36 | 0.34 | 0.30 | 0.02 | 0.32 | 0.32 | 0.32 | 0.34 | 0.18** | 0.17** | 0.15** | -0.03 | 0.03 | 0.04 | 0.02 | -0.13 |

This table presents a comprehensive comparison of four models (cVAE, GAMLSS, MFPR, and HBR) across various brain regions using four key performance metrics. These metrics were computed using predicted and actual data from the hold-out datasets to assess model generalisation. For Spearman's correlation coefficient, * indicates statistical significance with ($p < 0.05$), while ** indicates ($p < 0.01$). Abbreviations: cVAE = conditional Variational Autoencoder; GAMLSS = Generalised Additive Models for Location, Scale and Shape; MFPR = Multivariate Fractional Polynomial Regression; HBR = Hierarchical Bayesian Regression. See Table S1 for abbreviations of lobar and arterial regions.

**Table S6 Spearman's correlations between age and z-score across brain regions and models on the hold-out dataset.**

| | cVAE | | GAMLSS | | MFPR | | HBR | |
|---|---|---|---|---|---|---|---|---|
| | Coefficient | Corrected $p$-value | Coefficient | Corrected $p$-value | Coefficient | Corrected $p$-value | Coefficient | Corrected $p$-value |
| whole-brain | -0.022 | 0.384 | 0.033 | 0.208 | 0.095 | **<0.001** | 0.068 | **0.002** |
| periventricular | -0.019 | 0.463 | 0.024 | 0.376 | 0.098 | **<0.001** | 0.058 | **0.009** |
| deep white matter | -0.009 | 0.676 | 0.008 | 0.837 | 0.027 | 0.223 | 0.021 | 0.470 |
| lFrontal | -0.100 | **<0.001** | 0.002 | 0.994 | 0.206 | **<0.001** | 0.133 | **<0.001** |
| rFrontal | -0.049 | **0.025** | 0.001 | 0.994 | 0.174 | **<0.001** | 0.113 | **<0.001** |
| lTemporal | -0.121 | **<0.001** | -0.077 | **<0.001** | 0.075 | **<0.001** | 0.087 | **<0.001** |
| rTemporal | -0.113 | **<0.001** | -0.086 | **<0.001** | 0.023 | 0.283 | 0.064 | **0.004** |
| lParietal | -0.010 | 0.676 | 0.010 | 0.818 | 0.217 | **<0.001** | 0.152 | **<0.001** |
| rParietal | -0.073 | **0.001** | -0.028 | 0.303 | 0.148 | **<0.001** | 0.093 | **<0.001** |
| lOccipital | -0.061 | **0.004** | -0.031 | 0.228 | 0.050 | **0.017** | 0.036 | 0.139 |
| rOccipital | -0.087 | **<0.001** | -0.058 | **0.011** | 0.072 | **<0.001** | -0.006 | 0.900 |
| lCerebellum | **0.473** | **<0.001** | **0.470** | **<0.001** | -0.018 | 0.405 | 0.054 | **0.017** |
| rCerebellum | **0.513** | **<0.001** | 0.125 | **<0.001** | -0.057 | **0.007** | -0.004 | 0.922 |
| Brainstem | -0.018 | 0.463 | -0.033 | 0.208 | 0.061 | **0.004** | -0.001 | 0.987 |
| lAAH | -0.013 | 0.623 | -0.039 | 0.125 | -0.010 | 0.668 | 0.039 | 0.106 |
| rAAH | -0.039 | 0.079 | -0.049 | **0.044** | -0.049 | **0.019** | 0.002 | 0.987 |
| lMAH | 0.038 | 0.086 | 0.010 | 0.818 | 0.053 | **0.012** | 0.012 | 0.767 |
| rMAH | -0.043 | **0.049** | 0.002 | 0.994 | 0.055 | **0.009** | 0 | 0.996 |
| lAAML | 0.017 | 0.504 | 0.007 | 0.850 | 0.066 | **0.002** | 0.009 | 0.881 |

| | | | | | | | | |
|---|---|---|---|---|---|---|---|---|
| rAAML | -0.073 | **0.001** | -0.010 | 0.818 | 0.060 | **0.005** | -0.013 | 0.758 |
| lAAC | 0.090 | **<0.001** | 0.018 | 0.575 | 0.091 | **<0.001** | 0.006 | 0.900 |
| rAAC | 0.003 | 0.892 | 0 | 0.994 | 0.017 | 0.431 | -0.040 | 0.106 |
| lMALL | 0.011 | 0.676 | 0.037 | 0.152 | 0.113 | **<0.001** | 0.027 | 0.314 |
| rMALL | -0.061 | **0.004** | -0.001 | 0.994 | 0.056 | **0.008** | 0.024 | 0.392 |
| lPATMP | 0.101 | **<0.001** | 0.024 | 0.376 | 0.034 | 0.115 | 0.008 | 0.893 |
| rPATMP | -0.006 | 0.786 | 0.011 | 0.818 | 0.046 | **0.029** | 0.020 | 0.500 |
| lPAH | 0.020 | 0.448 | -0.043 | 0.090 | -0.001 | 0.957 | 0.031 | 0.227 |
| rPAH | -0.071 | **0.001** | -0.068 | **0.002** | -0.008 | 0.723 | 0.006 | 0.900 |
| lPAC | **-0.476** | **<0.001** | **-0.332** | **<0.001** | **0.305** | **<0.001** | 0.159 | **<0.001** |
| rPAC | **-0.547** | **<0.001** | **-0.386** | **<0.001** | **0.337** | **<0.001** | 0.139 | **<0.001** |

This table summarises Spearman's correlation coefficients and corrected *p*-values for the relationship between age and z-scores of white matter hyperintensity (WMH) volumes across various brain regions on the hold-out dataset. P-values were corrected for multiple comparisons using the False Discovery Rate (FDR) method. Abbreviations: cVAE = conditional Variational Autoencoder; GAMLSS = Generalised Additive Models for Location, Scale and Shape; MFPR = Multivariate Fractional Polynomial Regression; HBR = Hierarchical Bayesian Regression. See Table S1 for abbreviations of lobar and arterial regions.

**Table S7 Percentage of extreme deviations (%) on the hold-out dataset.**

| | cVAE | | GAMLSS | | MFPR | | HBR | |
|---|---|---|---|---|---|---|---|---|
| | Z > 2.58 | Z < -2.58 | Z > 2.58 | Z < -2.58 | Z > 2.58 | Z < -2.58 | Z > 2.58 | Z < -2.58 |
| whole-brain | 1.6 | 0.39 | 2.69 | 1.09 | 14.19 | 5.69 | 0.94 | 0.12 |
| periventricular | 1.91 | 0.66 | 2.61 | 1.09 | 14.54 | 6.08 | 1.01 | 0.19 |
| deep white matter | 0.27 | 1.05 | 1.36 | 3.04 | 11.54 | 6.82 | 0.23 | 1.48 |
| lFrontal | 0 | 5.46 | 0.08 | 6.98 | 0.12 | 9.35 | 0 | 6.55 |
| rFrontal | 0 | 5.96 | 0.04 | 7.52 | 0.23 | 11.03 | 0 | 8.81 |
| lTemporal | 0 | 0.27 | 0 | 3.16 | 0 | 4.56 | 0 | 10.37 |
| rTemporal | 0 | 0.04 | 0 | 2.18 | 0 | 3.66 | 0 | 5.14 |
| lParietal | 0 | 10.68 | 0.04 | 8.89 | 1.79 | 10.41 | 0 | 3.12 |
| rParietal | 0 | 9.43 | 0 | 8.38 | 2.73 | 10.48 | 0 | 0.62 |
| lOccipital | 0 | 2.22 | 0 | 1.21 | 0 | 5.77 | 0 | 1.48 |
| rOccipital | 0 | 0 | 0 | 10.52 | 0 | 17.58 | 0 | 1.91 |
| lCerebellum | 7.25 | 0 | 7.25 | 11.89 | 7.33 | 0.04 | 7.40 | 1.68 |
| rCerebellum | 6.43 | 0 | 6.43 | 5.65 | 6.39 | 0 | 6.47 | 0.51 |
| Brainstem | 0 | 11.85 | 0.08 | 3.31 | 2.26 | 11.96 | 0 | 0.74 |
| lAAH | 0 | 5.14 | 0.23 | 2.18 | 9.12 | 5.81 | 0 | 0.74 |
| rAAH | 0 | 3.12 | 0.27 | 3.08 | 13.02 | 4.44 | 0 | 0.51 |
| lMAH | 0.27 | 0.74 | 1.64 | 3.82 | 14.34 | 7.01 | 0.16 | 1.29 |
| rMAH | 0.43 | 1.29 | 1.71 | 2.96 | 11.93 | 7.37 | 0.27 | 1.17 |
| lAAML | 0.08 | 1.29 | 0.78 | 2.34 | 11.11 | 7.83 | 0.08 | 5.73 |
| rAAML | 0.04 | 1.09 | 0.82 | 2.10 | 10.60 | 6.70 | 0.12 | 2.57 |

| | | | | | | | | |
|---|---|---|---|---|---|---|---|---|
| lAAC | 0 | 0.51 | 0.47 | 2.26 | 8.03 | 7.09 | 0 | 0 |
| rAAC | 0 | 0.31 | 0.66 | 1.52 | 7.60 | 6.66 | 0 | 0 |
| lMALL | 0.35 | 0.66 | 1.36 | 3.20 | 12.47 | 7.33 | 0.27 | 0 |
| rMALL | 0.74 | 0.27 | 2.53 | 2.81 | 13.87 | 6.43 | 0.47 | 0 |
| lPATMP | 0.04 | 1.01 | 0.94 | 6.00 | 8.26 | 9.39 | 0.19 | 0 |
| rPATMP | 0.04 | 0.86 | 1.05 | 3.08 | 9.66 | 8.96 | 0.19 | 0 |
| lPAH | 0.04 | 5.73 | 0.08 | 0 | 6.63 | 6.51 | 0 | 0 |
| rPAH | 0.04 | 2.61 | 0.12 | 0 | 6.59 | 5.42 | 0 | 0 |
| lPAC | 15.08 | 0 | 8.77 | 0 | 14.93 | 0 | 19.91 | 0 |
| rPAC | 11.54 | 0 | 8.65 | 0 | 11.30 | 0 | 12.47 | 0 |

This table presents the percentage of samples with extreme deviations (z-score > 2.58 or z-score < -2.58) across models on the hold-out dataset. Abbreviations: cVAE = conditional Variational Autoencoder; GAMLSS = Generalised Additive Models for Location, Scale and Shape; MFPR = Multivariate Fractional Polynomial Regression; HBR = Hierarchical Bayesian Regression. See Table S1 for abbreviations of lobar and arterial regions.

**Table S8 Spearman's correlations between WMH volume Z-scores and hypertension levels across brain regions and models.**

| | cVAE | | GAMLSS | | MFPR | | HBR | |
|---|---|---|---|---|---|---|---|---|
| | Coefficient | Corrected $p$-value | Coefficient | Corrected $p$-value | Coefficient | Corrected $p$-value | Coefficient | Corrected $p$-value |
| whole-brain | 0.077 | **<0.001** | 0.082 | **<0.001** | 0.089 | **<0.001** | 0.093 | **<0.001** |
| periventricular | 0.079 | **<0.001** | 0.079 | **<0.001** | 0.089 | **<0.001** | 0.090 | **<0.001** |
| deep white matter | 0.058 | **<0.001** | 0.067 | **<0.001** | 0.068 | **<0.001** | 0.070 | **<0.001** |
| lFrontal | 0.016 | **0.026** | 0.044 | **<0.001** | 0.092 | **<0.001** | 0.072 | **<0.001** |
| rFrontal | 0.037 | **<0.001** | 0.059 | **<0.001** | 0.093 | **<0.001** | 0.080 | **<0.001** |
| lTemporal | 0.016 | **0.022** | 0.034 | **<0.001** | 0.078 | **<0.001** | 0.069 | **<0.001** |
| rTemporal | 0.013 | 0.070 | 0.022 | **<0.001** | 0.072 | **<0.001** | 0.065 | **<0.001** |
| lParietal | 0.050 | **<0.001** | 0.053 | **<0.001** | 0.086 | **<0.001** | 0.090 | **<0.001** |
| rParietal | 0.038 | **<0.001** | 0.052 | **<0.001** | 0.081 | **<0.001** | 0.086 | **<0.001** |
| lOccipital | 0.029 | **<0.001** | 0.028 | **<0.001** | 0.056 | **<0.001** | 0.045 | **<0.001** |
| rOccipital | 0.025 | **<0.001** | 0.039 | **<0.001** | 0.063 | **<0.001** | 0.049 | **<0.001** |
| lCerebellum | 0.093 | **<0.001** | 0.062 | **<0.001** | -0.015 | **0.036** | 0.044 | **<0.001** |
| rCerebellum | 0.103 | **<0.001** | -0.003 | 0.697 | -0.004 | 0.552 | -0.002 | 0.746 |
| Brainstem | 0.011 | 0.113 | 0 | 0.970 | 0.030 | **<0.001** | 0.013 | 0.057 |
| lAAH | 0.045 | **<0.001** | 0.046 | **<0.001** | 0.048 | **<0.001** | 0.061 | **<0.001** |
| rAAH | 0.043 | **<0.001** | 0.047 | **<0.001** | 0.047 | **<0.001** | 0.057 | **<0.001** |
| lMAH | 0.070 | **<0.001** | 0.062 | **<0.001** | 0.066 | **<0.001** | 0.064 | **<0.001** |
| rMAH | 0.065 | **<0.001** | 0.071 | **<0.001** | 0.077 | **<0.001** | 0.073 | **<0.001** |
| lAAML | 0.065 | **<0.001** | 0.061 | **<0.001** | 0.069 | **<0.001** | 0.063 | **<0.001** |
| rAAML | 0.036 | **<0.001** | 0.047 | **<0.001** | 0.059 | **<0.001** | 0.049 | **<0.001** |

| | | | | | | | | |
|---|---|---|---|---|---|---|---|---|
| lAAC | 0.091 | **<0.001** | 0.070 | **<0.001** | 0.084 | **<0.001** | 0.070 | **<0.001** |
| rAAC | 0.065 | **<0.001** | 0.058 | **<0.001** | 0.060 | **<0.001** | 0.050 | **<0.001** |
| lMALL | 0.079 | **<0.001** | 0.076 | **<0.001** | 0.087 | **<0.001** | 0.075 | **<0.001** |
| rMALL | 0.072 | **<0.001** | 0.070 | **<0.001** | 0.078 | **<0.001** | 0.079 | **<0.001** |
| lPATMP | 0.046 | **<0.001** | 0.026 | **<0.001** | 0.027 | **<0.001** | 0.025 | **<0.001** |
| rPATMP | 0.028 | **<0.001** | 0.026 | **<0.001** | 0.034 | **<0.001** | 0.028 | **<0.001** |
| lPAH | 0.066 | **<0.001** | 0.043 | **<0.001** | 0.050 | **<0.001** | 0.061 | **<0.001** |
| rPAH | 0.048 | **<0.001** | 0.047 | **<0.001** | 0.058 | **<0.001** | 0.063 | **<0.001** |
| lPAC | -0.063 | **<0.001** | -0.062 | **<0.001** | 0.054 | **<0.001** | 0.062 | **<0.001** |
| rPAC | -0.091 | **<0.001** | -0.074 | **<0.001** | 0.077 | **<0.001** | 0.074 | **<0.001** |

This table presents the results of Spearman's correlation analyses between White Matter Hyperintensity (WMH) volume Z-scores and hypertension levels (0-3) for different brain regions across four models. P-values were corrected for multiple comparisons using the False Discovery Rate (FDR) method. Abbreviations: cVAE = conditional Variational Autoencoder; GAMLSS = Generalised Additive Models for Location, Scale and Shape; MFPR = Multivariate Fractional Polynomial Regression; HBR = Hierarchical Bayesian Regression. See Table S1 for abbreviations of lobar and arterial regions.

**Table S9 Percentage of extreme deviations (%) in the evaluation dataset.**

| | cVAE | | | | GAMLSS | | | | MFPR | | | | HBR | | | |
|---|---|---|---|---|---|---|---|---|---|---|---|---|---|---|---|---|
| **Hypertension Level** | **0** | **1** | **2** | **3** | **0** | **1** | **2** | **3** | **0** | **1** | **2** | **3** | **0** | **1** | **2** | **3** |
| whole-brain | 1.60 | 3.44 | 4.48 | 6.96 | 2.69 | 4.46 | 5.17 | 7.59 | 14.19 | 18.75 | 21.2 | 27.89 | 0.94 | 2.53 | 3.20 | 5.61 |
| periventricular | 1.91 | 4.11 | 5.64 | 8.52 | 2.61 | 4.11 | 5.16 | 7.52 | 14.54 | 19.18 | 21.55 | 27.89 | 1.01 | 2.37 | 3.25 | 5.96 |
| deep white matter | 0.27 | 0.28 | 0.29 | 0.28 | 1.36 | 2.74 | 2.96 | 4.68 | 11.54 | 14.37 | 16.65 | 21.22 | 0.23 | 0.54 | 0.48 | 0.71 |
| lFrontal | 0 | 0 | 0 | 0 | 0.08 | 0.26 | 0.32 | 0.64 | 0.12 | 0.96 | 1.30 | 2.70 | 0 | 0 | 0 | 0 |
| rFrontal | 0 | 0 | 0 | 0 | 0.04 | 0.12 | 0.16 | 0.43 | 0.23 | 0.99 | 1.35 | 2.34 | 0 | 0 | 0 | 0 |
| lTemporal | 0 | 0 | 0 | 0 | 0 | 0.01 | 0 | 0 | 0 | 0.12 | 0.21 | 0.35 | 0 | 0 | 0 | 0 |
| rTemporal | 0 | 0 | 0 | 0 | 0 | 0 | 0 | 0 | 0 | 0.04 | 0.02 | 0 | 0 | 0 | 0 | 0 |
| lParietal | 0 | 0 | 0 | 0 | 0.04 | 0.26 | 0.48 | 0.92 | 1.79 | 4.34 | 5.61 | 8.16 | 0 | 0 | 0 | 0 |
| rParietal | 0 | 0 | 0 | 0 | 0 | 0.11 | 0.21 | 0.57 | 2.73 | 5.50 | 7.50 | 12.14 | 0 | 0 | 0 | 0 |
| lOccipital | 0 | 0 | 0 | 0 | 0 | 0 | 0 | 0 | 0 | 0 | 0 | 0 | 0 | 0 | 0 | 0 |
| rOccipital | 0 | 0 | 0 | 0 | 0 | 0 | 0 | 0 | 0 | 0 | 0.02 | 0 | 0 | 0 | 0 | 0 |
| lCerebellum | 7.25 | 7.53 | 8.07 | 7.59 | 7.25 | 7.53 | 8.07 | 7.59 | 7.33 | 7.55 | 8.12 | 7.74 | 7.40 | 7.69 | 8.58 | 8.30 |
| rCerebellum | 6.43 | 7.27 | 6.89 | 6.10 | 6.43 | 7.27 | 6.89 | 6.10 | 6.39 | 7.26 | 6.86 | 6.10 | 6.47 | 7.36 | 7.01 | 6.32 |
| Brainstem | 0 | 0 | 0 | 0 | 0.08 | 0.13 | 0.24 | 0.14 | 2.26 | 4.32 | 5.43 | 6.46 | 0 | 0 | 0 | 0 |
| lAAH | 0 | 0 | 0 | 0 | 0.23 | 0.38 | 0.50 | 0.99 | 9.12 | 12.23 | 13.03 | 17.74 | 0 | 0 | 0 | 0 |
| rAAH | 0 | 0 | 0 | 0 | 0.27 | 0.75 | 0.80 | 1.42 | 13.02 | 16.12 | 17.63 | 21.58 | 0 | 0 | 0 | 0 |
| lMAH | 0.27 | 0.51 | 0.59 | 0.99 | 1.64 | 2.49 | 2.83 | 3.97 | 14.34 | 17.15 | 18.25 | 23.78 | 0.16 | 0.24 | 0.24 | 0.43 |
| rMAH | 0.43 | 0.83 | 1.03 | 1.42 | 1.71 | 2.91 | 3.52 | 4.90 | 11.93 | 15.78 | 17.31 | 23.07 | 0.27 | 0.54 | 0.58 | 0.92 |
| lAAML | 0.08 | 0.05 | 0.08 | 0.21 | 0.78 | 1.12 | 1.61 | 3.26 | 11.11 | 14.66 | 16.46 | 20.37 | 0.08 | 0.18 | 0.21 | 0.57 |
| rAAML | 0.04 | 0.07 | 0.08 | 0.14 | 0.82 | 1.34 | 1.75 | 2.56 | 10.6 | 13.69 | 15.17 | 20.65 | 0.12 | 0.17 | 0.24 | 0.64 |

| | | | | | | | | | | | | | | | | |
|---|---|---|---|---|---|---|---|---|---|---|---|---|---|---|---|---|
| lAAC | 0 | 0 | 0 | 0.07 | 0.47 | 0.67 | 0.76 | 1.63 | 8.03 | 12.04 | 14.4 | 17.46 | 0 | 0 | 0 | 0.07 |
| rAAC | 0 | 0 | 0 | 0 | 0.66 | 0.48 | 0.43 | 0.78 | 7.60 | 10.49 | 11.79 | 14.62 | 0 | 0 | 0 | 0 |
| lMALL | 0.35 | 0.43 | 0.02 | 1.35 | 1.36 | 2.07 | 2.36 | 4.19 | 12.47 | 17.43 | 19.33 | 25.27 | 0.27 | 0.34 | 0.29 | 0.92 |
| rMALL | 0.74 | 1.52 | 0.61 | 3.55 | 2.53 | 2.87 | 3.17 | 5.25 | 13.87 | 17.34 | 19.56 | 24.77 | 0.47 | 1.12 | 1.51 | 3.05 |
| lPATMP | 0.04 | 0.09 | 1.94 | 0.21 | 0.94 | 1.23 | 1.32 | 2.06 | 8.26 | 8.41 | 9.27 | 10.15 | 0.19 | 0.28 | 0.35 | 0.35 |
| rPATMP | 0.04 | 0.10 | 0.16 | 0.28 | 1.05 | 1.67 | 1.54 | 2.91 | 9.66 | 10.66 | 11.67 | 14.83 | 0.19 | 0.40 | 0.34 | 0.78 |
| lPAH | 0.04 | 0 | 0.13 | 0.07 | 0.08 | 0.25 | 0.27 | 0.85 | 6.63 | 8.48 | 9.64 | 13.77 | 0 | 0 | 0 | 0 |
| rPAH | 0.04 | 0.02 | 0.02 | 0 | 0.12 | 0.37 | 0.42 | 1.42 | 6.59 | 10.75 | 12.6 | 18.10 | 0 | 0 | 0 | 0 |
| lPAC | 15.08 | 16.89 | 16.94 | 20.01 | 8.77 | 7.61 | 6.72 | 7.10 | 14.93 | 16.51 | 16.33 | 17.39 | 19.91 | 25.26 | 28.15 | 35.70 |
| rPAC | 11.54 | 15.55 | 17.90 | 22.78 | 8.65 | 10.19 | 10.19 | 11.43 | 11.30 | 14.24 | 15.81 | 18.81 | 12.47 | 16.71 | 19.54 | 25.05 |

This table presents the percentage of samples with extreme deviations (z-score > 2.58) across hypertension levels and models in the evaluation dataset. Abbreviations: cVAE = conditional Variational Autoencoder; GAMLSS = Generalised Additive Models for Location, Scale and Shape; MFPR = Multivariate Fractional Polynomial Regression; HBR = Hierarchical Bayesian Regression. See Table S1 for abbreviations of lobar and arterial regions.

#### Supplementary Figures

##### Figure S1 Distribution of Whole-Brain White Matter Hyperintensity (WMH) Volume: Original vs. Normalised

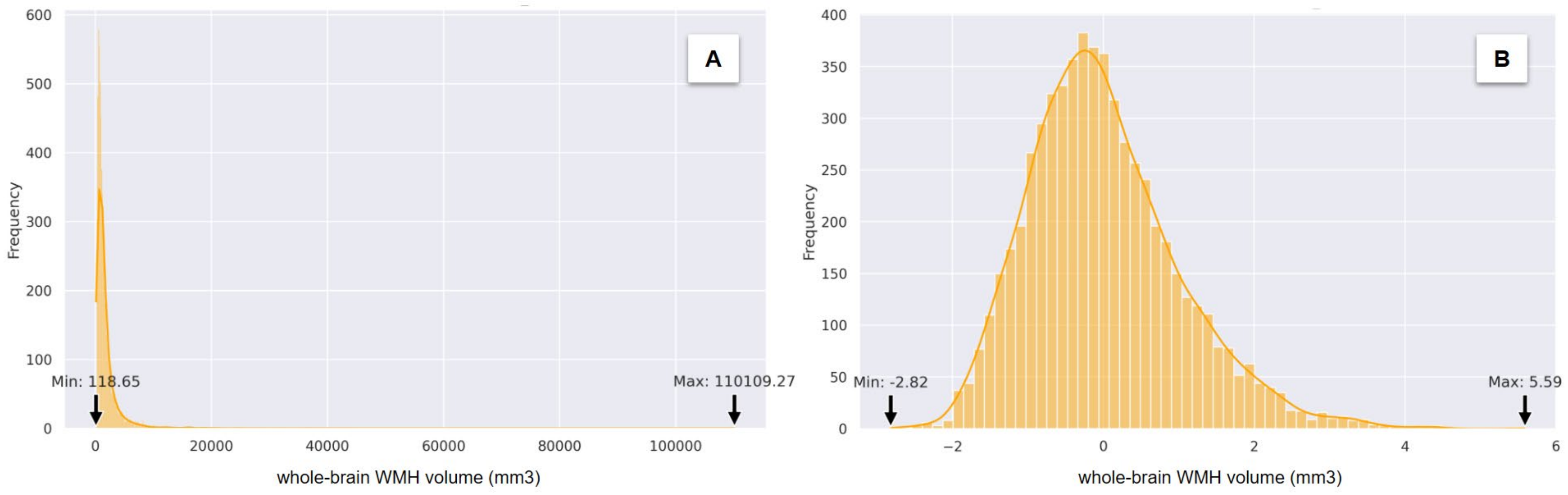

Panel A shows the distribution of whole-brain WMH volumes in their original scale (in mm³). Panel B presents the log-transformed and standardised distribution of the same data.

**Figure S2 Heatmaps of Spearman Correlations Between Covariates and Brain Region-Specific Measures in the Training Dataset.**

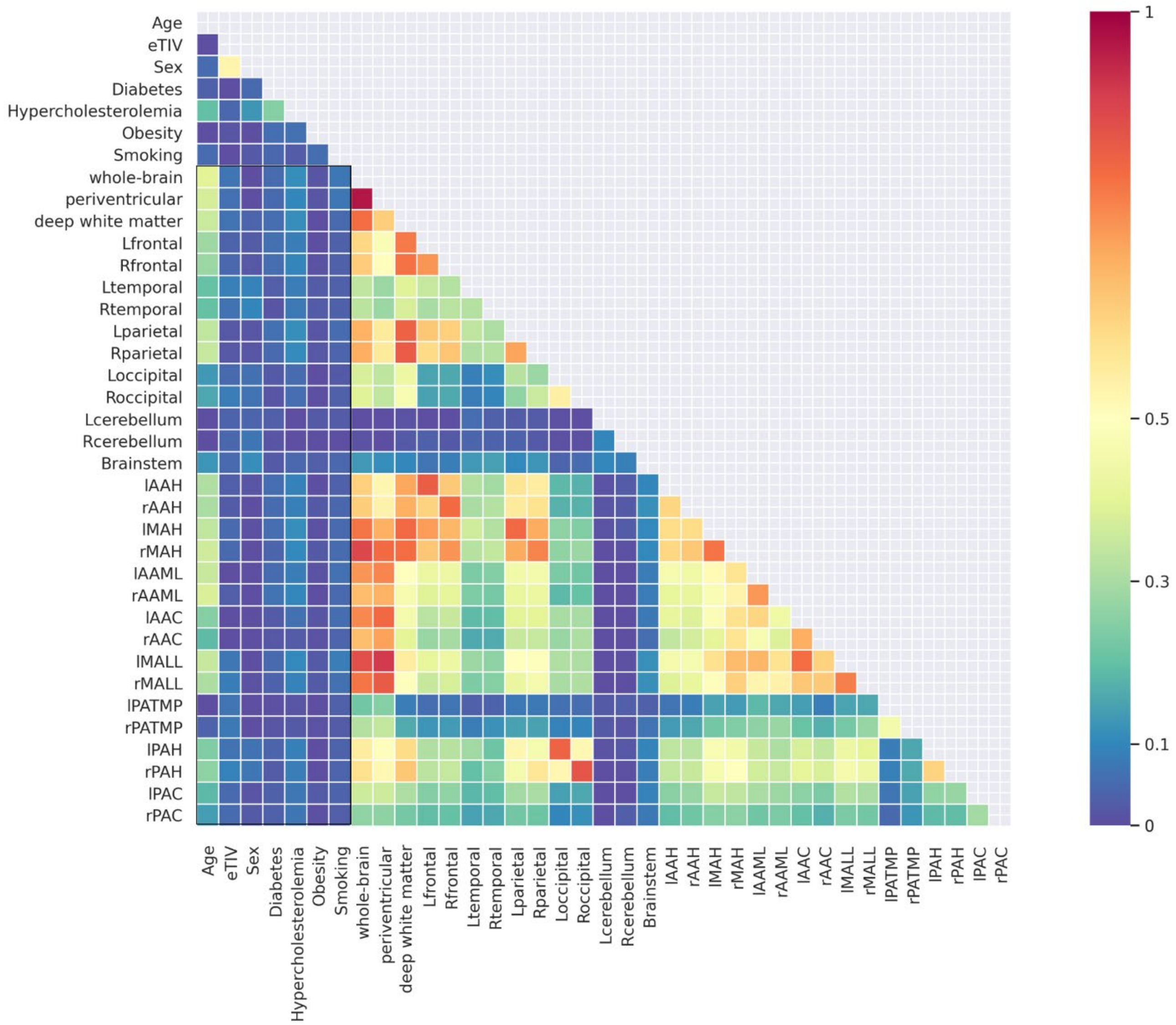

**Figure S3 Training and Validation Loss over Epochs.**

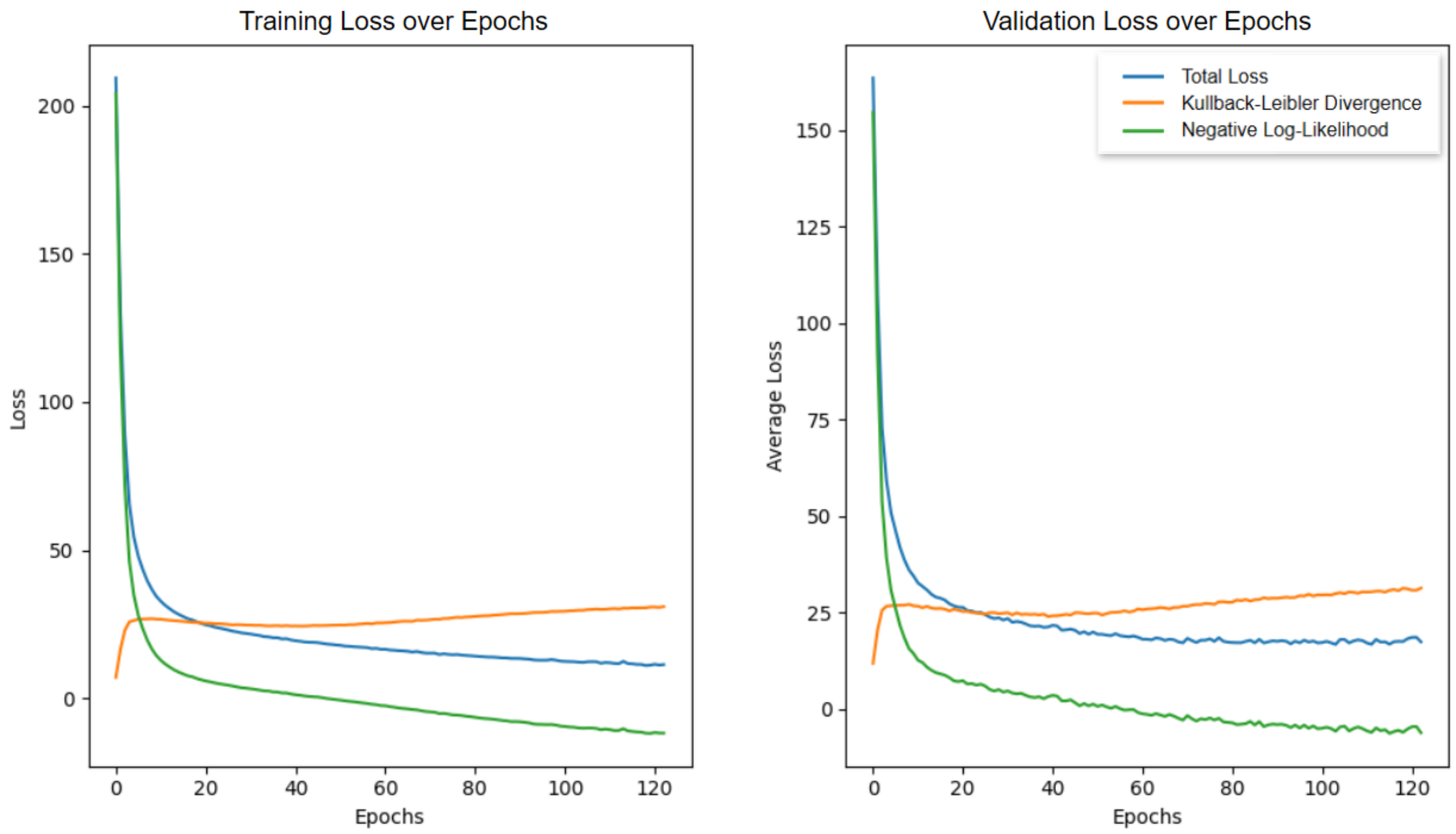

This plot shows the training (left) and validation (right) loss components during the training of a conditional Variational Autoencoder (cVAE) model across epochs. An early stopping algorithm was applied to prevent overfitting, which triggered at epoch 123.

**Figure S4 Bland-Altman Plots and Distribution of Quantile Loss at 90th Percentile for cVAE vs Other Models.**

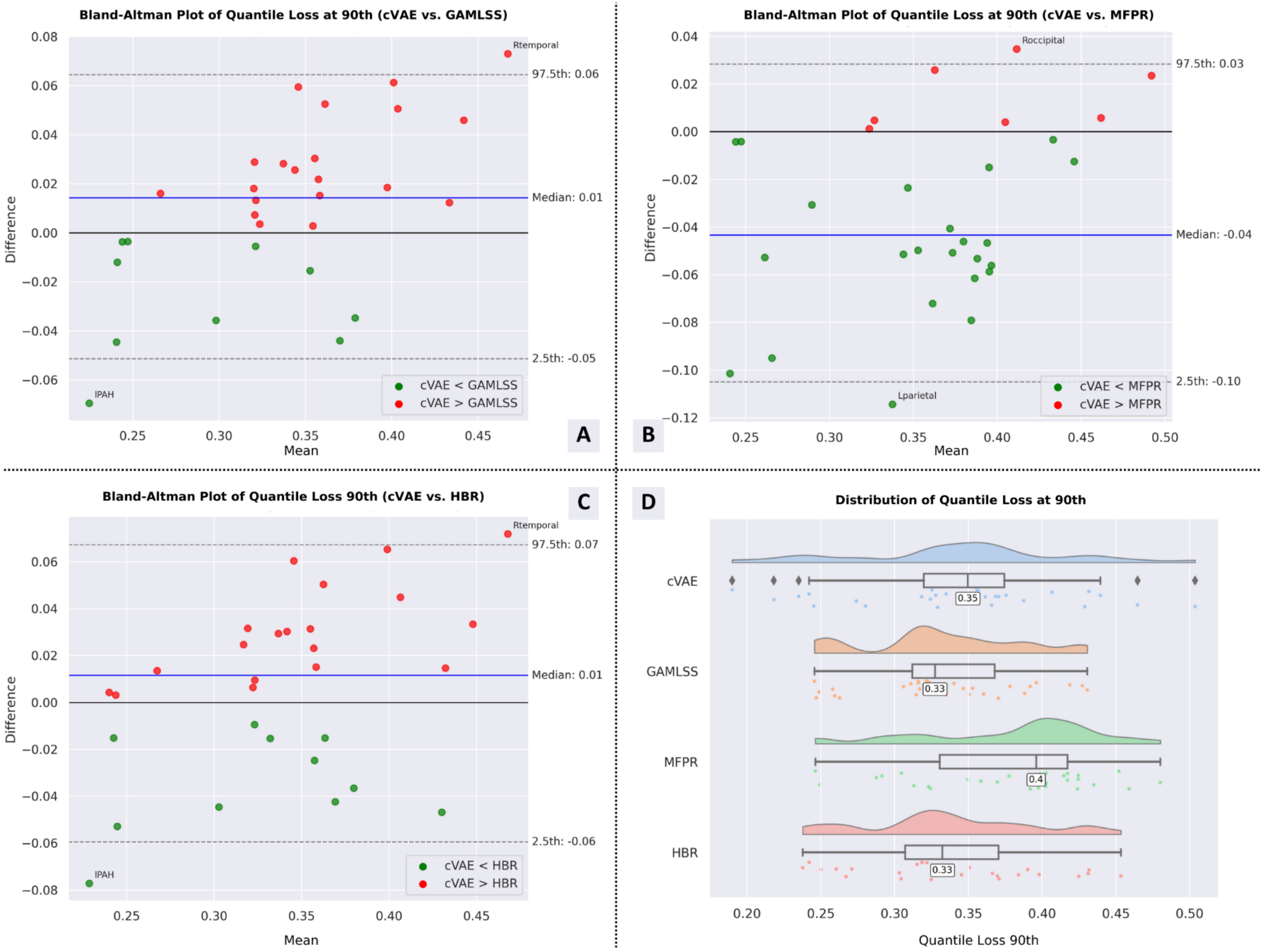

**(A-C)** Bland-Altman plots comparing the Quantile Loss at the 90th percentile of the conditional Variational Autoencoder (cVAE) model against the Generalised Additive Models for Location, Scale, and Shape (GAMLSS) (A), Multivariate Fractional Polynomial Regression (MFPR) (B), and Hierarchical Bayesian Regression (HBR) (C) models. Each point represents a different brain region. Green points indicate regions where cVAE exhibits lower Quantile Loss at the 90th percentile compared to the corresponding model, while red points show regions where the other model performs better.

**(D)** Distribution of Quantile Loss at the 90th percentile across all four models (cVAE, GAMLSS, MFPR, and HBR), visualising the variability and central tendency for each model's performance. This metric is computed on the hold-out dataset.

**Figure S5 Bland-Altman Plots and Distribution of Spearman Correlation for cVAE vs Other Models.**

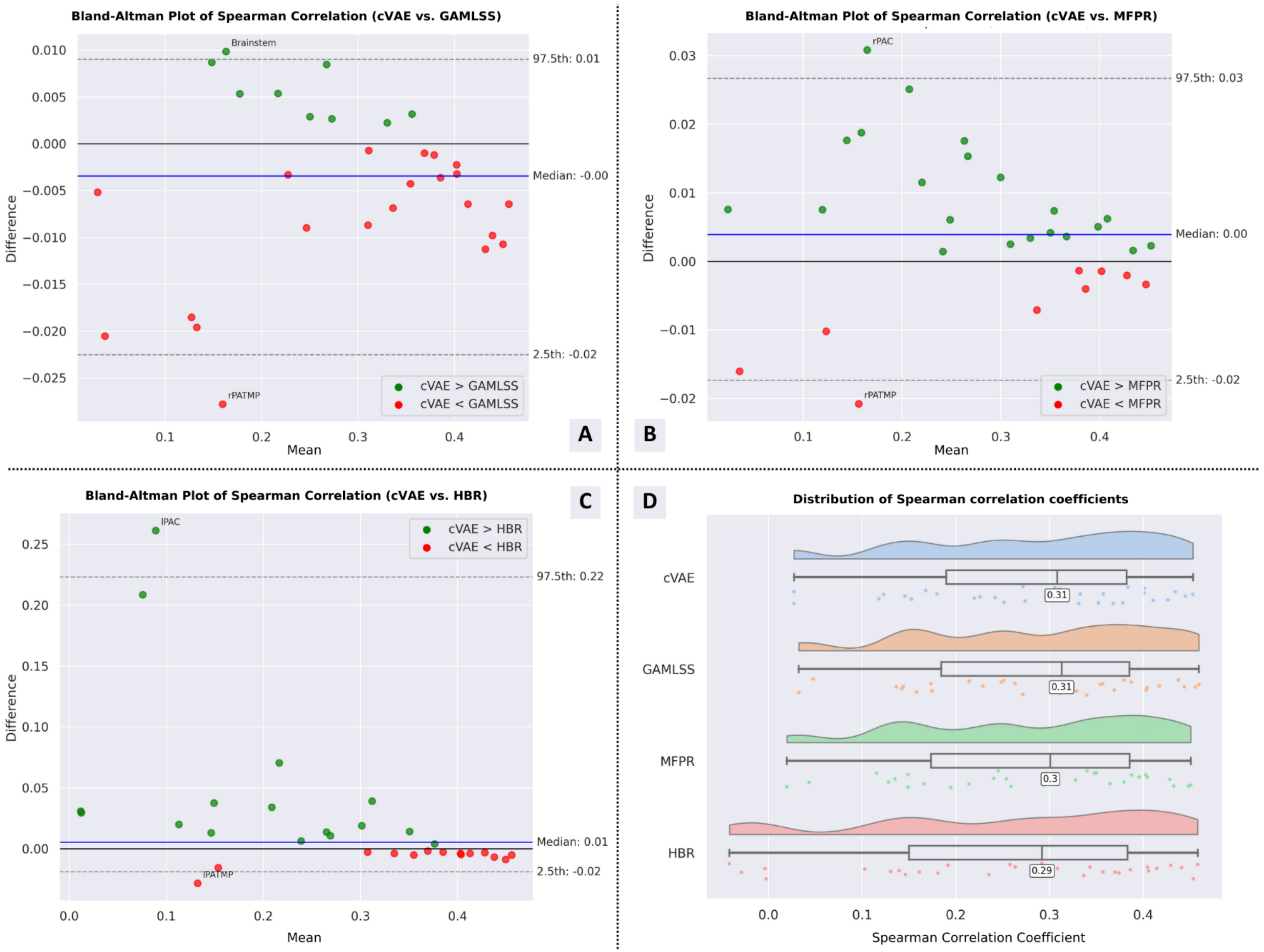

**(A-C)** Bland-Altman plots comparing the Spearman Correlation of the conditional Variational Autoencoder (cVAE) model against the Generalised Additive Models for Location, Scale, and Shape (GAMLSS) (A), Multivariate Fractional Polynomial Regression (MFPR) (B), and Hierarchical Bayesian Regression (HBR) (C) models. Each point represents a different brain region. Green points indicate regions where cVAE exhibits higher Explained Variance than the corresponding model, while red points show regions where the other model performs better.

**(D)** Distribution of Explained Variance across all four models (cVAE, GAMLSS, MFPR, and HBR), visualising the variation and central tendency for each model's performance. This metric is computed on the hold-out dataset.

**Figure S6 Bland-Altman Plots and Distribution of Explained Variance for cVAE vs Other Models.**

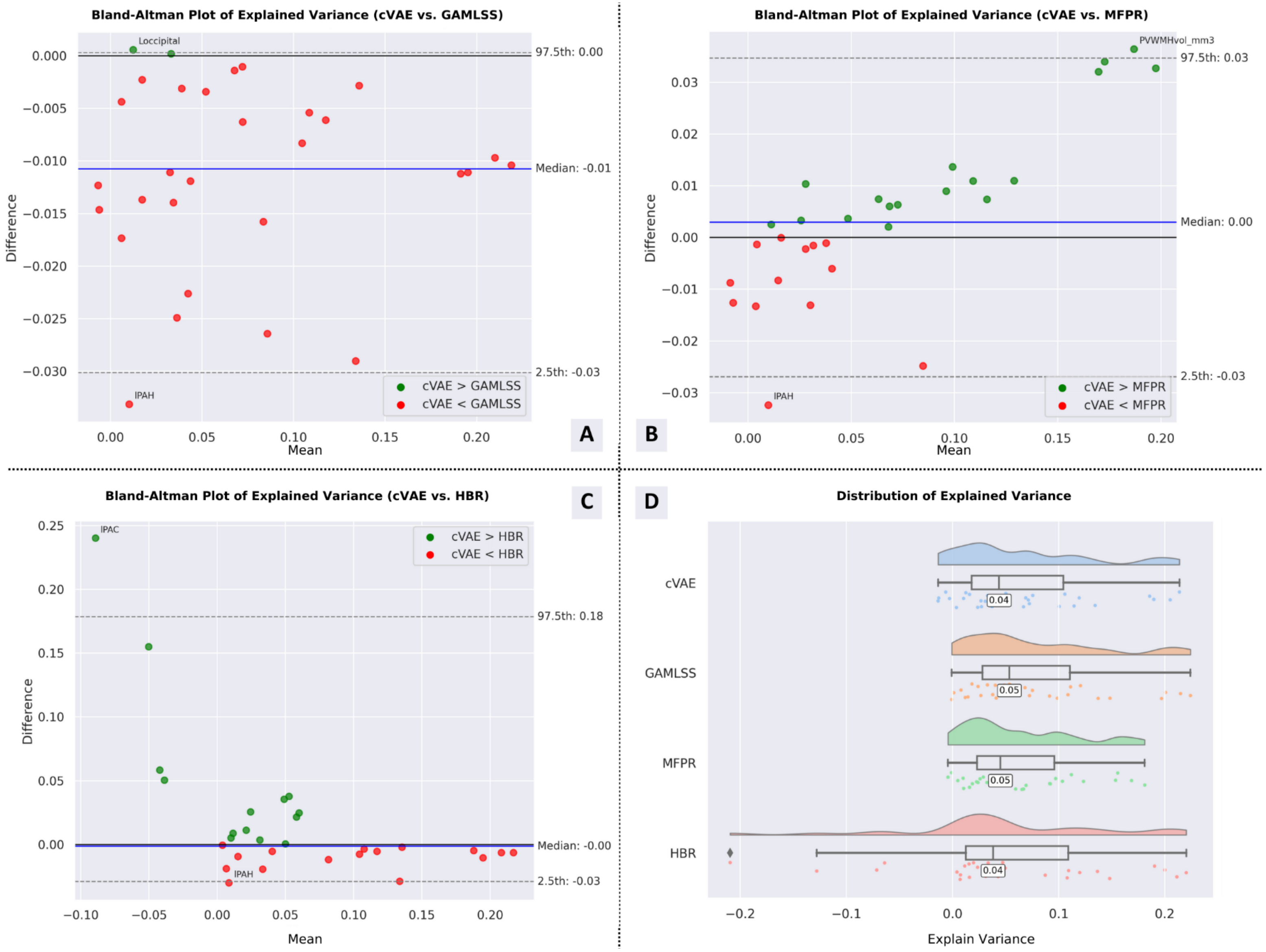

**(A-C)** Bland-Altman plots comparing the Explained Variance of the conditional Variational Autoencoder (cVAE) model against the Generalised Additive Models for Location, Scale, and Shape (GAMLSS) (A), Multivariate Fractional Polynomial Regression (MFPR) (B), and Hierarchical Bayesian Regression (HBR) (C) models. Each point represents a different brain region. Green points indicate regions where cVAE exhibits higher Explained Variance than the corresponding model, while red points show regions where the other model performs better.

**(D)** Distribution of Explained Variance across all four models (cVAE, GAMLSS, MFPR, and HBR), visualising the variation and central tendency for each model's performance. This metric is computed on the hold-out dataset.

**Figure S7 Bland-Altman Plots and Distribution of Median Z-score for cVAE vs Other Models.**

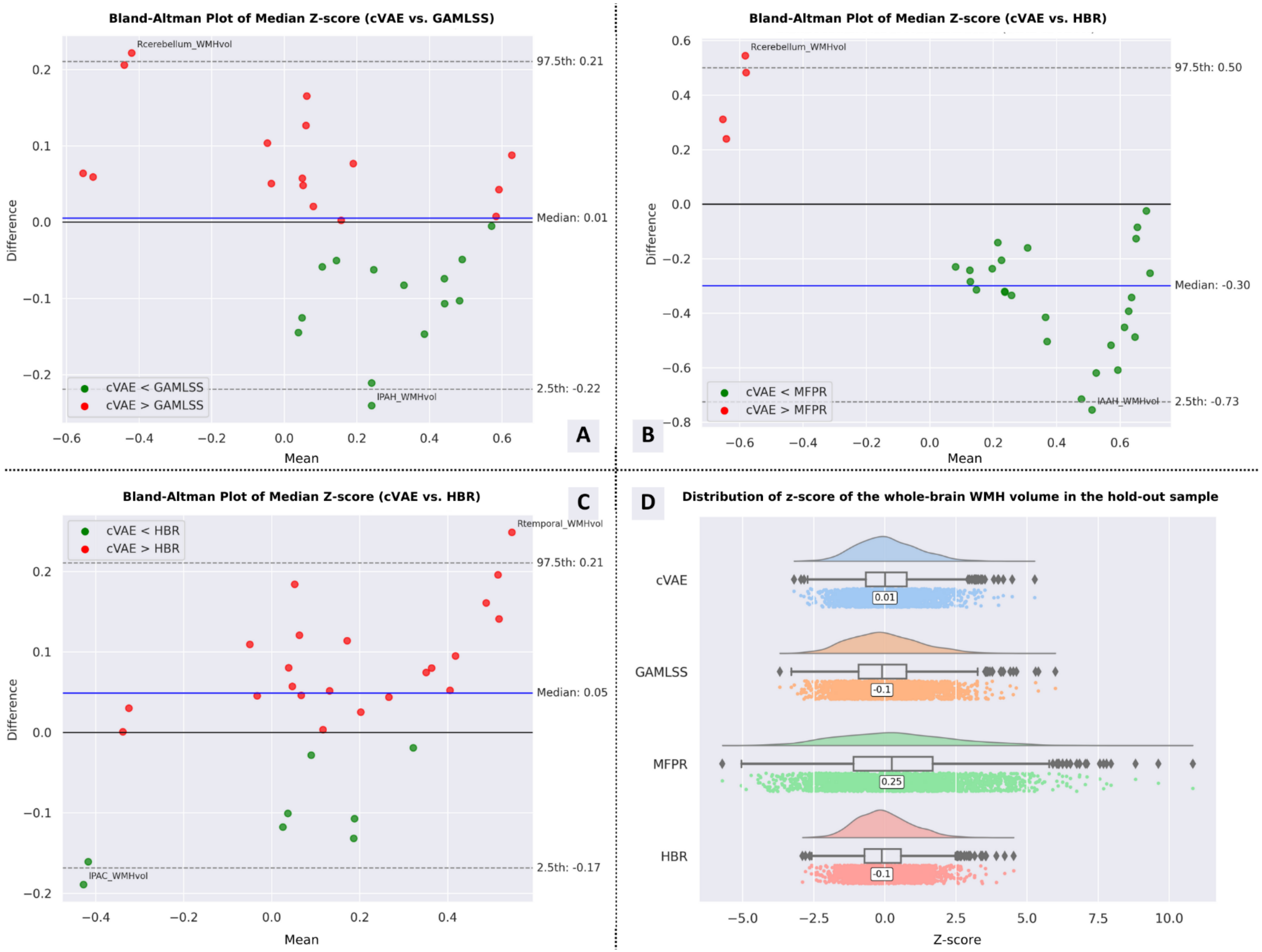

**(A-C)** Bland-Altman plots comparing the median Z-scores of cVAE against GAMLSS (A), MFPR (B), and HBR (C) models on the hold-out dataset. Green points indicate where cVAE performs better (lower median Z-scores) than the compared model, while red points show the opposite. Each point represents a different brain region (see Table S1).

**(D)** Distribution of Z-scores for the whole-brain white matter hyperintensity (WMH) volume across all models (cVAE, GAMLSS, MFPR, and HBR). This metric is computed on the hold-out dataset.

**Figure S8 Spearman Correlation between Z-score and Hypertension levels.**

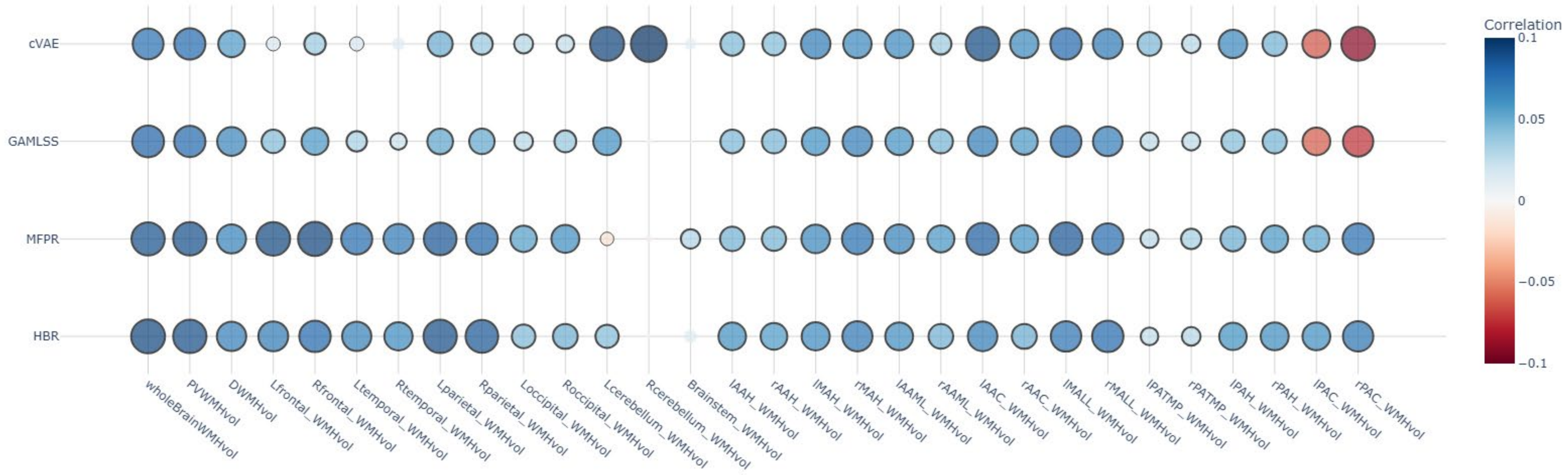

This bubble chart illustrates the Spearman correlation between z-scores and hypertension levels (0-3) in the evaluation dataset, comparing different models (cVAE, GAMLSS, MFPR, HBR) across various brain regions. Bubble size and colour intensity represent correlation strength, with darker blue indicating stronger positive correlations and darker red showing stronger negative correlations. Bubbles with borders denote statistically significant correlations (False Discovery Rate (FDR)-corrected p-value < 0.05). Abbreviations: cVAE = conditional Variational Autoencoder; GAMLSS = Generalised Additive Models for Location, Scale and Shape; MFPR = Multivariate Fractional Polynomial Regression; HBR = Hierarchical Bayesian Regression; WMH = White Matter Hyperintensity. See Table S1 for brain region-specific measure abbreviations.
